# Temporal scaling in developmental gene networks by epigenetic timing control

**DOI:** 10.1101/752170

**Authors:** Phuc H.B. Nguyen, Nicholas A. Pease, Kenneth K.H. Ng, Blythe Irwin, Hao Yuan Kueh

## Abstract

During development, progenitors follow defined temporal schedules for differentiation, to form organs and body plans with precise sizes and proportions. Across diverse contexts, these developmental schedules are encoded by autonomous timekeeping mechanisms in single cells. These autonomous timers not only operate robustly over many cell generations, but can also operate at different speeds in different species, enabling proportional scaling of temporal schedules and population sizes. By combining mathematical modeling with live-cell measurements, we elucidate the mechanism of a polycomb-based epigenetic timer, that delays activation of the T-cell commitment regulator *Bcl11b* to facilitate progenitor expansion. This mechanism generates activation delays that are independent of cell cycle duration, and are tunably controlled by transcription factors and epigenetic modifiers. When incorporated into regulatory gene networks, this epigenetic timer enables progenitors to set scalable temporal schedules for flexible size control. These findings illuminate how evolution may set and adjust developmental speed in multicellular organisms.

## INTRODUCTION

How are sizes and proportions of organs and body plans set during multicellular development, and how can they be adjusted through evolutionary changes? The developmental mechanisms controlling size and form must not only facilitate flexible adjustment of individual multicellular assemblies, but must also enable proportionally scaled changes in the sizes of different assemblies, to flexibly vary total organism size. For over a century, it has been recognized that the timing of developmental events in the vertebrate embryo is a central determinant of size and form (De Beer, 1940; Gould, 1977; Haeckel, 1866; Huxley, 1942). Because progenitors continually grow and divide during development, the timing at which they differentiate determines their degree of expansion and their final population size; consequently, changes in developmental timing can engender variation in size or form. Changes in the timing of individual lineage specification events can give rise to innovations in shape or form (Alberch et al., 1979; Gould, 1977; Huxley, 1932), whereas changes to the overall speed of development can lead to proportionally scaled changes in organ and organism sizes (Bonner, 1965; Calder, 1984).

As embryonic development is not coupled to an external clock, the temporal schedules for lineage specification must be controlled by mechanisms intrinsic to the embryo itself (Ebisuya and Briscoe, 2018). These developmental schedules are generally the product of complex processes unfolding in space and time in embryo; however, across diverse contexts, there is growing evidence that these schedules may largely be set by autonomous timekeeping mechanisms operating in single progenitors (Figure 1A) (Burton et al., 1999; Gao et al., 1997; Heinzel et al., 2017; Otani et al., 2016; Rosello-Diez et al., 2014; Saiz-Lopez et al., 2015). For example, during cerebral cortex development, neural progenitors follow a defined temporal schedule to generate different cortical neurons, where they generate inner layer neurons before outer layer neurons. Strikingly, this schedule is recapitulated in progenitors cultured *in vitro* in the presence of constant inductive signals, even in the absence of an intact three-dimensional tissue environment (Eiraku et al., 2008; Gaspard et al., 2008). Furthermore, while the order of differentiation remains the same in progenitors from different species, the temporal schedule itself is expanded or contracted in an autonomous manner, with total durations ranging from one week in mouse progenitors, to months in human progenitors (van den Ameele et al., 2014; Barry et al., 2017; Espuny-Camacho et al., 2013; Otani et al., 2016). Such scalability in temporal schedules over long timescales enables progenitors to vary output numbers of different cell types while keeping their proportions constant, and may thus underlie the variability in organ and body plan sizes across evolution (Figure 1).

**Figure 1.**
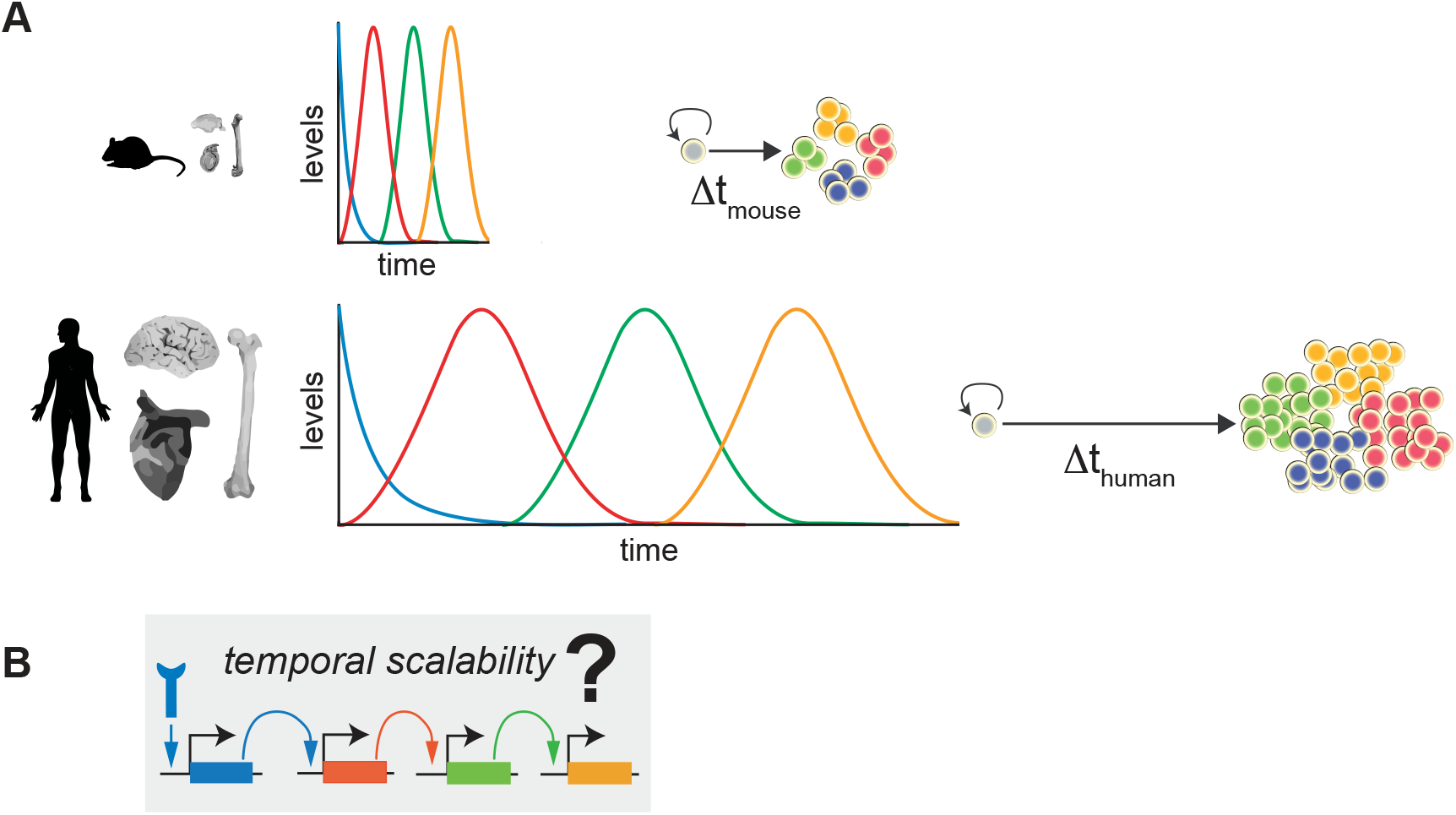
Temporal scalability in development. (A) In different developmental systems, dividing progenitors maintain cell-autonomous schedules for lineage specification. These temporal schedules can be compressed or extended in time for proportional scaling of organ and organism size. (B) Developmental schedules are executed by conserved developmental gene networks; however, it remains unlear what molecular processes allow gene networks to set autonomous, temporally-scalable schedules over develompental timescales.

Conserved networks of regulatory genes control cell type specification during development, and ultimately dictate when or whether progenitor cells turn on lineage-specifying genes in response to signals (Alon, 2007; Briscoe and Small, 2015; Davidson, 2010; Georgescu et al., 2008; Medina et al., 2005). However, it is unknown how developmental gene networks can generate cell-autonomous time schedules that are robust, yet scalable by evolution (Figure 1B). Specifically, there are two major issues: First, it is unclear how the developmental gene networks can robustly operate over timescales spanning many cell generations. In gene networks built from classical biophysical models of gene regulation (Ackers et al., 1982; Alon, 2007; Bintu et al., 2005; Bolouri and Davidson, 2002), network dynamics are determined by the mRNA and protein half-lives, which are constrained by cell division and dilution; consequently, it is difficult to robustly generate gene activation delays with timescales longer than one cell generation (Levine and Elowitz, 2014). Secondly, it is unclear how temporal schedules set by developmental gene networks can be coordinately expanded or contracted by evolutionary variation. Changes in the timing of individual developmental events can occur through *cis*-regulatory element mutations at individual regulatory gene loci, as is indeed observed (Frankel et al., 2011; Gérard et al., 1997; Khan et al., 2011; Simeonov et al., 2017; Walters et al., 1995; Weintraub, 1988); however, it is unclear how coordinated, scaled changes in activation timing could within an entire regulatory network, particularly between related species where network components are highly conserved.

Epigenetic mechanisms, involving the polycomb repressive system and its cognate histone H3K27 trimethylation (H3K27me3) mark, are important for differentiation in all multicellular eukaryotes (Aloia et al., 2013; Xiao and Wagner, 2015), and may play central roles in developmental timing control. H3K27me3 modifications are broadly found at lineage-specifying gene loci in stem and progenitor cells (Boyer et al., 2006; Lee et al., 2006), and are removed in response to signals to enable regulatory gene activation and differentiation. While cells differentiate with a time delay after sensing developmental signals, H3K27me3 loss is often assumed to rapidly follow the binding of transcription factors (Kaneko et al., 2014; Rank et al., 2002; Riising et al., 2014), whose slow accumulation is thought to set the pace for downstream gene activation. However, other studies have found that the process of H3K27me3 removal may itself be slow (Berry et al., 2017; Kaikkonen et al., 2013; Mayran et al., 2018), and may generate long time delays prior to gene activation and cell differentiation (Angel et al., 2011; Bintu et al., 2016), even when upstream transcription factors are fully present. An ability to generate delays, together with broad action at many gene loci, could enable H3K27me3 and polycomb regulators to act coordinately at many loci in regulatory gene networks to control developmental speed and population size expansion. However, the dynamics and biophysical mechanisms by which these factors control regulatory gene activation remain unclear, because it has been difficult to resolve these processes in living cells.

To gain insights into these questions, we investigate an epigenetic mechanism controlling the activation timing of *Bcl11b*, a T-cell lineage commitment regulator that turns on with an extended delay to enable early progenitor expansion (Figure 2A). After entering the thymus, hematopoietic progenitor cells turn on *Bcl11b* to commit to the T-cell lineage (Hosokawa et al., 2018; Ikawa et al., 2010; Li et al., 2010a, 2010b). *Bcl11b* activation requires Notch signals in the thymus, along with multiple transcription factors that are up-regulated by Notch signaling (García-Ojeda et al., 2013; Germar et al., 2011; Li et al., 2010b; Weber et al., 2011). However, while all these factors are induced shortly after thymic entry, *Bcl11b* activation and T-cell lineage commitment occurs only ∼5-10 days later, during which cells expand by approximately a thousand-fold (Manesso et al., 2013; Porritt et al., 2003). To determine whether this time delay may be the result of epigenetic mechanisms acting on individual *Bcl11b* loci, we generated a reporter mouse, where two *Bcl11b* loci were tagged with distinguishable fluorescent protein expression reporters (Ng et al., 2018). By separately following the activation dynamics of two *Bcl11b* loci in single progenitors from these mice, we found evidence that a slow, stochastic epigenetic event, occurring on the *Bcl11b* locus, generates a multi-day time delay in *Bcl11b* activation and T-cell commitment (Figure 2A). This finding shows that epigenetic mechanisms can act in *cis* to generate time delays in gene activation and fate commitment, and establishes a unique model for studying these mechanisms in single, living cells.

**Figure 2.**
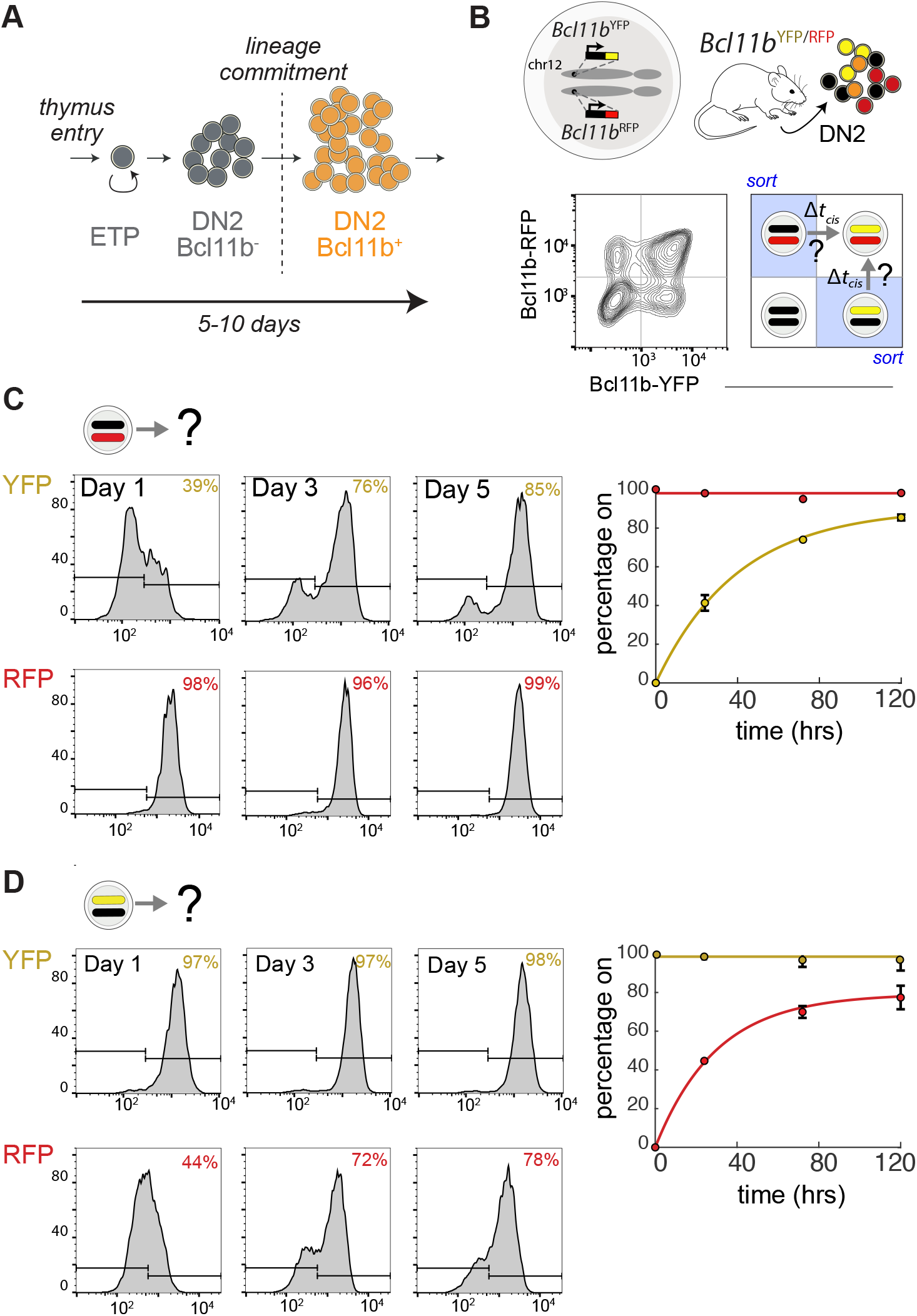
An epigenetic timing mechanism, acting on the *Bcl11b* locus, delays gene activation and T-cell lineage commitment. (A) Schematic of early T-cell development. Early cell expansion is enabled by delayed activation of the T-cell commitment regulator Bcl11b. (B) Dual-allelic *Bcl11b* reporter mouse (top), along with flow cytometry plot showing levels of the two *Bcl11b* reporters in bone-marrow derived DN2 progenitors (bottom left), along with strategy to purify monoallelic progenitors for live analysis of epigenetic timing step. (C-D) DN2 monoallelic progenitors were purified by cell sorting, cultured on OP9-DL1 with 5 ng/mL IL-7 and Flt3L, and analyzed by flow cytometry. Mean values and 95% confidence intervals are plotted for *n* = 3 different batches of bone marrow. Curves represent fits to the equation f(x) = F(1-e^-kt^) (k = 0.025 hrs^-1^ +/-0.005 for Bcl11b-YFP and k = 0.034 hrs^-1^ +/-0.009 for Bcl11b-RFP).

In this study, we first elucidate the mechanism of the epigenetic timer controlling *Bcl11b* activation. To do so, we combine quantitative, single-cell measurements in this dual-reporter system with analysis of candidate biophysical mechanisms using mathematical modeling. Next, by modeling developmental gene networks built from these epigenetic timers, we propose a solution for how temporal schedules for development can be scaled to vary size while maintaining proportions. We find that this epigenetic timing mechanism involves delayed, all-or-none removal of a repressive H3K27me3 domain from the gene locus, and can set delays that are robust over many cell generations, yet are tunable by activities of histone-modifying enzymes. The delays set by this epigenetic timer are unaffected by changes in cell cycle duration, enabling independent control of proliferation and developmental timing in developing progenitors. When incorporated as a building block of developmental gene networks, this epigenetic timer enables progenitors to set schedules for differentiation that are flexibly expandible or compressible in time, by tuning of the strength of epigenetic repression. These findings establish a biophysical basis for understanding dynamic gene activation control by epigenetic mechanisms, and reveal insights into how developmental speed and organism size are controlled in development and evolution.

## RESULTS

The dual-allelic *Bcl11b* reporter system provides a unique, powerful tool to resolve epigenetic mechanisms controlling gene activation timing in living cells. To study the *cis*-epigenetic event controlling *Bcl11b* activation timing in isolation from other *trans-* events, we can analyze *Bcl11b* locus activation dynamics in progenitors that already have one *Bcl11b* allele active, and therefore must contain all *trans*-factors necessary for expression (Figure 2B). Using fluorescence-activated cell sorting, we purified monoallelic expressing *Bcl11b* DN2 progenitors from dual-allelic reporter mice (YFP^+^RFP^-^ or YFP^-^RFP^+^), and analyzed activation of the silent allele by co-culture with OP9-DL1 cells, an *in vitro* system that recapitulates all early transitions in T-cell development. In agreement with previous work (Ng et al., 2018), the inactive *Bcl11b* allele turned on in an all-or-none manner, with slow onset timing over the course of five days (Figure 2C-D). Furthermore, activation kinetics were similar for both YFP and RFP alleles (k = 0.025 +/-0.005 hrs^-1^ for YFP allele and k = 0.034 +/-0.009 hrs^-1^ for RFP allele), and were well described by a single exponential curve, consistent with activation being controlled by a single stochastic event occurring with equal likelihood at the two loci, with a timescale of days.

### Developmental timing control by H3K27me3 mark removal

Polycomb associated histone modification H3K27me3 is highly enriched at silent *Bcl11b* loci in hematopoietic stem and progenitor cells but not in committed T-cells, where *Bcl11b* is expressed (Zhang et al., 2012); thus, its removal could underlie the *cis-*epigenetic event controlling *Bcl11b* activation timing. To test this possibility, we first determined whether H3K27me3 marks are removed from the *Bcl11b* locus at the same time it turns on. To pinpoint when H3K27me3 loss occurs relative to locus activation, we measured H3K27me3 levels in three populations having different numbers of active *Bcl11b* alleles: hematopoietic progenitor cells from bone marrow, which have both *Bcl11b* alleles inactive; monoallelic *Bcl11b* expressing DN2 progenitors, which have one active and one inactive *Bcl11b* allele; and biallelic *Bcl11b* expressing DN2 progenitors, which have both *Bcl11b* alleles active. We employed CUT&RUN, a novel nuclease-based method for mapping DNA-protein complexes that can be combined with spike-in controls to provide quantitative readouts of H3K27me3 genomic abundance comparable across samples (Skene et al., 2018). If H3K27me3 marks are removed concurrently with *Bcl11b* activation, but not any sooner or later, we would expect H3K27me3 levels in monoallelic DN2 progenitors to fall to approximately half of the initial levels found in HSPCs and to approximately zero in biallelic DN2 progenitors. Indeed, H3K27me3 levels at the *Bcl11b* locus decreased across these populations in a manner consistent these predictions. In bone marrow progenitors, where both *Bcl11b* alleles are inactive, there was an abundance of H3K27me3 across 5’ end of *Bcl11b* (Figure 3A, yellow shaded region). These broad H3K27me3 peaks were roughly halved in monoallelic *Bcl11b* expressing cells, and were almost completely absent in *Bcl11b* biallelic cells (Figure 3A, 0.5 and 0.1 for monoallelic and biallelic *Bcl11b* expressing progenitors, respectively). Thus, H3K27me3 repressive mark removal from the *Bcl11b* locus occurs concurrently with gene activation.

**Figure 3:**
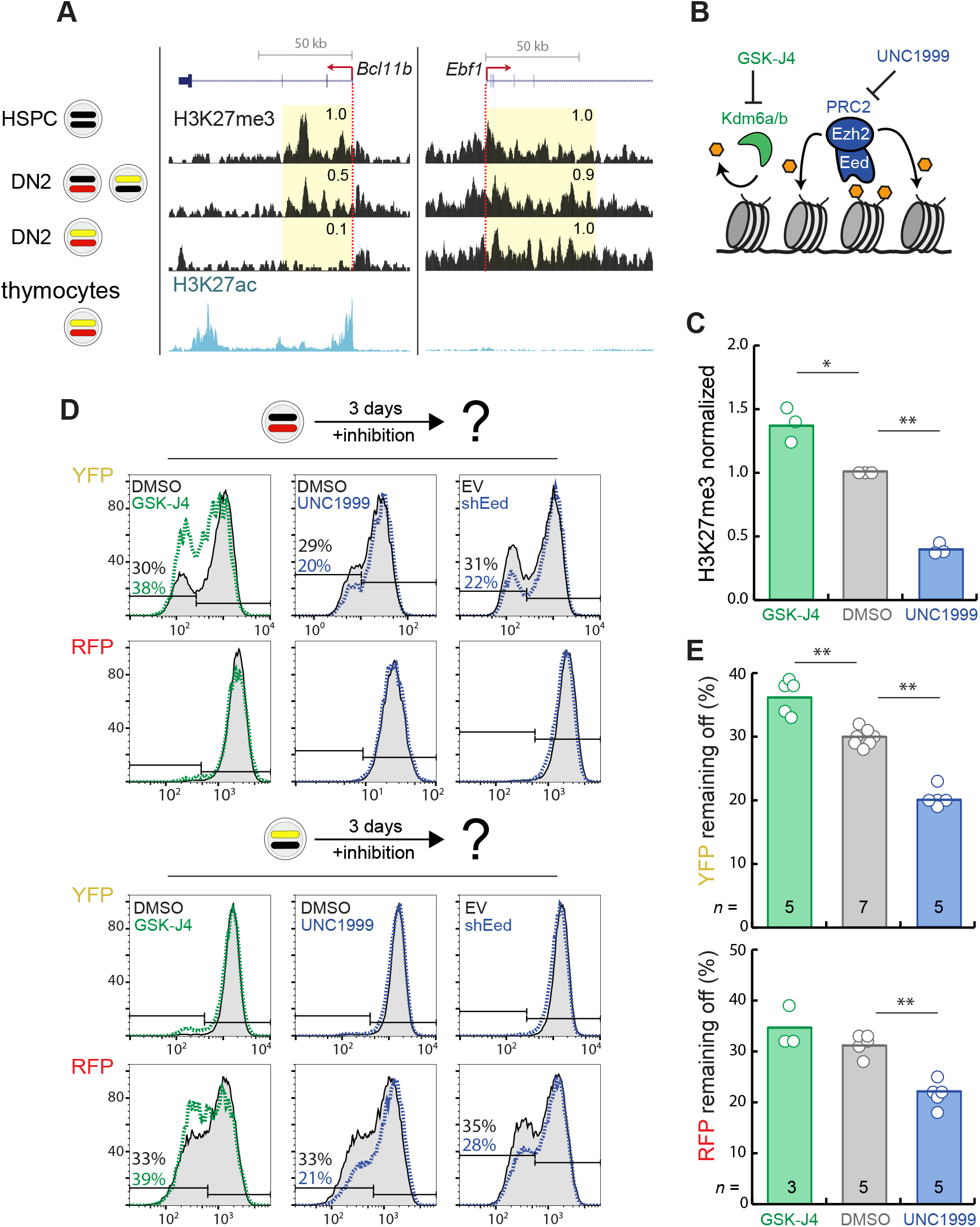
*Bcl11b* activation timing is tunably controlled by removal of repressive H3K27 trimethylation. (A) H3K27me3 distributions were profiled by CUT&RUN in Lin^-^ bone marrow progenitors (HSPCs), as well as sorted DN2 mono-allellic and bi-allelic cells. Genome browser tracks show H3K27me3 densities at *Bcl11b*, as well as at *Ebf1*, a B-cell regulator that is repressed in these cells. Relative densities in shaded areas are shown. H3K27ac levels in thymocytes, obtained from a previous study (Davis et al., 2018; Accession: ENCSR000CCH), are shown. Data are representative of two independent replicates (C) DN2 monoallelic progenitors treated with the indicated inhibitors were sorted for H3K27me3 CUT&RUN qPCR at the *Bcl11b* promoter. Mean values are presented with *n* = 3 experimental replicates (*p < 0.05, ** p < 0.01, paired t-test, two-tailed. (D-E) Sorted DN2 monoallelic cells were recultured with the indicated perturbations, and analyzed after 3 days. Means values are presented *n* = independent experiments (*p < 0.05, **p < 0.01, paired t-test, two-tailed).

H3K27me3 mark loss may control the timing of *Bcl11b* activation; alternatively, it may simply be a consequence of gene activation, due to clearance of methylated nucleosomes by active transcription (Hosogane et al., 2016; Kraushaar et al., 2013). To determine whether H3K27me3 loss plays a causal role in controlling *Bcl11b* activation timing, we purified monoallelic *Bcl11b* expressing DN2 progenitors, co-cultured them on OP9-DL1 cells for three days with small molecule inhibitors targeting H3K27me3-modifying enzymes, and analyzed the resultant effects on activation of the silent *Bcl11b* locus (Figure 3B). These inhibitors, which target either the PRC2 methyltransferase subunit Ezh2 (UNC1999, 2.5 μM) or the H3K27 demethylases Kdm6a/b (GSK-J4, 2.5 μM), resulted in a ∼40% increase and ∼60% decrease, respectively, in H3K27me3 abundance at the *Bcl11b* promoter of *Bcl11b* monoallelic cells (Figure 3C), indicating that they indeed modulate H3K27me3 levels at inactive *Bcl11b* loci.

In the absence of any inhibitors, the majority of progenitors activated the silent *Bcl11b* allele after three days, leaving about ∼30% remaining inactive, as expected (Figure 3D-E). Kdm6 demethylase inhibition increased the fraction of progenitors with remaining inactive *Bcl11b* alleles (Figure 3D, 3E, from 30% to 36% YFP inactive; p = 0.007). Impeded *Bcl11b* activation reflects a specific consequence of Kdm6a/b inhibition, as similar effects were observed with IOX-1, another inhibitor of Kdm6a/b, but not with Daminozide, a broad demethylase inhibitor that does not target Kdm6a/b (Figure S1A). Conversely, Ezh2 inhibition decreased the fraction of progenitors with remaining inactive alleles (Figure 3E from 30% to 20% RFP inactive; p = 0.001; 30% to 21%YFP inactive; p = 0.007). Similar effects were observed using structurally unrelated Ezh2 inhibitors GSK-126 and GSK-343 (Figure S1A) as well as small hairpin RNAs to knock down the expression of Eed, another essential PRC2 subunit (Figure 3D, S1B). We note that all the H3K27me3 perturbations tested had no effect on *Bcl11b* expression levels after activation (Fig. 3D), consistent with a specific role for this modification system in the control of gene activation timing. Taken together, these results indicate that H3K27me3 removal from the *Bcl11b* locus plays a causal role in controlling *Bcl11b* activation timing, and further show that the time constant for this activation event is tunably set by the balance between PRC2 and Kdm6a/b demethylase activities at the gene locus.

### A mathematical model of epigenetic timer function

In a number of systems, loss of H3K27me3 occurs through passive dilution of these repressive marks with DNA replication (Coleman and Struhl, 2017; Sun et al., 2014), a mechanism that could potentially account for the long activation delays spanning many cell generations observed (Ng et al., 2018; Fig. 1). However, the involvement of both PRC2 and Kdm6a/b demethylases in setting the timing for *Bcl11b* activation, as observed (Fig. 3), argues against such a passive H3K27me3 dilution mechanism, and instead suggests an active mechanism, whereby the activities of opposing H3K27me3-modifying enzymes at the *Bcl11b* locus set the timing of H3K27me3 elimination and gene activation. However, it is unclear what kinds of biophysical mechanisms may underlie the timing control at the *Bcl11b* locus.

Any mechanism to explain the *Bcl11b* epigenetic timer must account for its observed emergent properties, namely: 1) its ability to robustly set time delays that span multiple cell generations; 2) its stochastic nature; and 3) its tunability by histone modifying enzyme activities. To identify mechanisms that could potentially account for these properties, we analyzed a series of candidate biophysical mechanisms using mathematical modeling. H3K27me3 can bind PRC2 at an allosteric site to stimulate its methyltransferase activity (Margueron et al., 2009), a cooperative mechanism that is thought to maintain repressive H3K27me3 marks across cell division. Therefore, we first considered a simple model, where individual nucleosomes in a one-dimensional array undergo H3K27me3 methylation catalyzed by the presence of nearby methylated nucleosomes, demethylation, as well as H3K27me3 loss due to random nucleosome segregation during DNA replication (Figure 4A; see Mathematical Appendix) (Coleman and Struhl, 2017). Similar models have been shown to support multi-stable histone modification states that are heritable across cell division (Dodd et al., 2007; Zhang et al., 2014). In agreement, we found that single loci could switch from a H3K27 methylated, repressed state to a demethylated state with stochastic delays spanning multiple cell divisions (Figure 4B). However, in our simulations, averaged activation timing was extremely sensitive to H3K27me3 methylation levels in the silent state, with sensitivity coefficients far exceeding those derived from experimental data (Figure 3C, 3E; methylation only model *s* ∼14 vs. experimental value *s* ∼ 0.3-0.6), such that minor changes (∼10%) causing drastic timing changes (∼300 fold) (Figure 4C). This extreme sensitivity was ubiquitous across different parameter regimes (data not shown) and was also found in other studies (Dodd et al., 2007; Zhang et al., 2014), indicating that it represents a general feature of such switching models. By analyzing this system using a transition state theory framework (see Mathematical Appendix and Figure S10A), we found that switching times scale exponentially with methylation or demethylation rates, thus explaining the observed extreme sensitivity. Thus, models that consider histone modification dynamics alone are inconsistent with the tunable control of activation by H3K27me3-modifying enzymes observed experimentally (Figure 3D, E).

**Figure 4:**
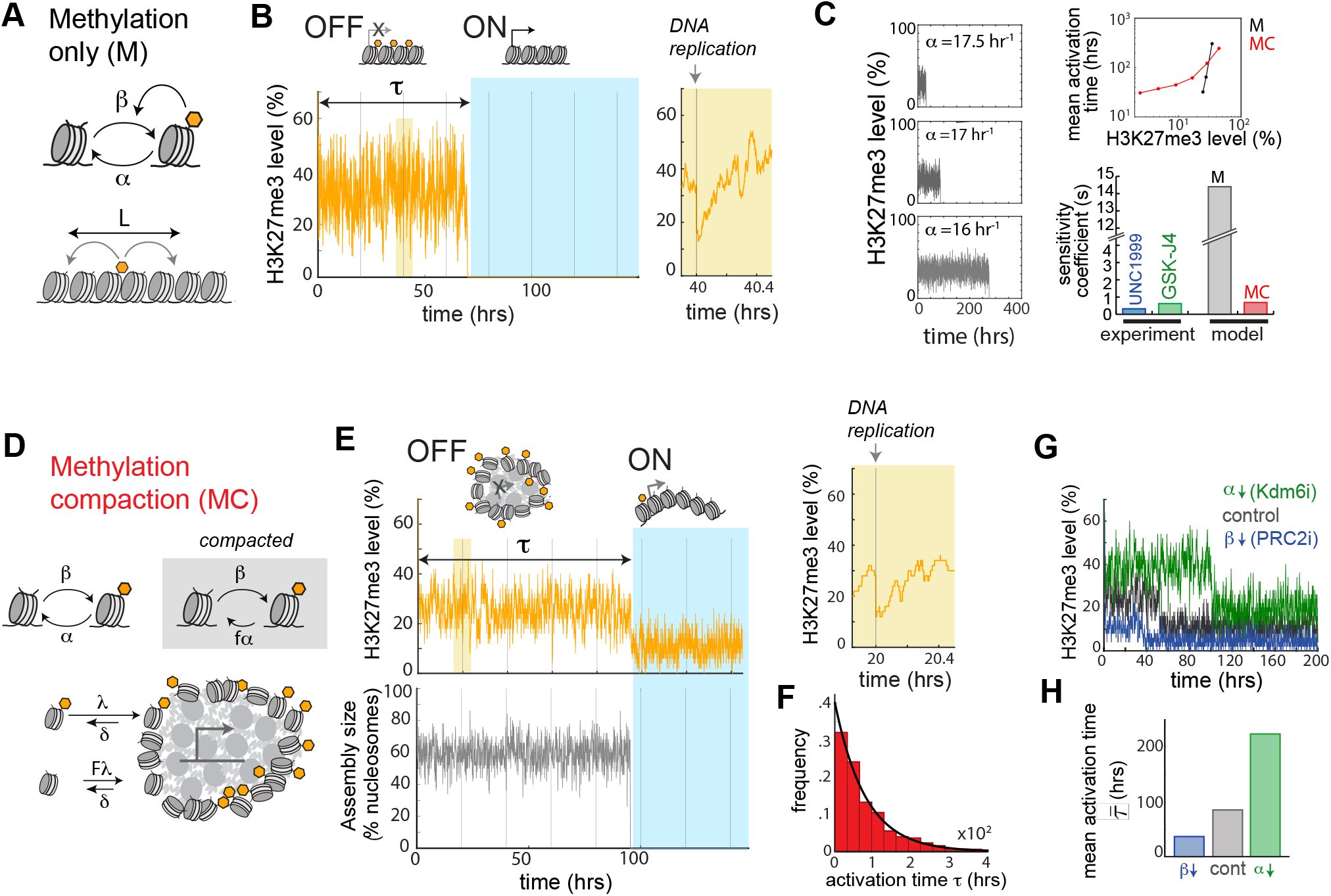
A methylation-compaction mechanism enables robust, tunable epigenetic timing control. Two candidate biophysical mechanisms for the epigenetic timer were analyzed using mathematical modeling, a methylation-only mechanism (A-C), and a methylation-compaction mechanism (D-H). (A) Methylation only model, along with (B) fraction H3K27me3 nucleosomes over time from a stochastic simulation, with inset showing H3K27me3 loss and recovery after DNA replication. (C) Sensitivity analysis of the methylation-only model (gray) and methylation-compaction model (red), showing H3K27me3 time traces for indicated parameters in the methylation model (left), mean activation time versus averaged H3K27me3 levels (top right), and averaged sensitivity coefficient s = *d*ln(y)/*d*ln(x) for the relationship between these two variables, both in models and experiments (blue, green, calculated from Fig. 3C,E). (D) The methylation-compaction model, along with (E) time traces of the fraction of H3K27me3-marked nucleosomes (top left), and the fraction of nucleosomes in a compacted assembly (bottom left). Inset (top right) shows H3K27me3 recovery after cell division. (F) Histogram shows distribution of activation times, along with fit to the exponential function y = e^-kt^ with k = 0.012 hr^-1^. H3K27me3 time traces (top) (G) and bar charts (H) show impact of simulated PRC2 or Kdm6 inhibition on initial H3K27me3 levels and mean activation times. Results represent average of 200 simulations.

H3K27me3 marks repress gene expression by promoting the association of nucleosomes to form condensed, polymerase-inaccessible assemblies. For instance, H3K27me3 recruits Polycomb Repressive Complex (PRC1), which can self-interact to form compacted, phase-separated chromatin domains (Plys et al., 2018; Tatavosian et al., 2018). Therefore, we considered a second model, where H3K27me3 levels do not determine gene synthesis rates *per se*, but influence chromatin compaction dynamics to modulate promoter accessibility and gene transcription (Figure 4D). In this methylation-compaction (MC) model, nucleosomes adopt methylated and demethylated states, as before; in addition, they can also bind or dissociate from a compacted assembly, with methylated nucleosomes binding with stronger affinity compared to demethylated nucleosomes. We do not explicitly model the spatial conformation of the compacted assembly; instead we adopt a chemical kinetics framework, approximating the assembly to be sphere with a minimum nucleus size, following established approaches to describe self-assembly of polymeric assemblies (Howard, 2001; Mitchison, 1992) (see Mathematical Appendix for details).

From simulations, we found that the system maintained a steady compacted, methylated state for multiple cell divisions before transitioning to uncompacted, demethylated state (Figure 4E). As with the methylation-only model, the timing of this all-or-none switch is well described by a first-order stochastic process, with a fixed probability of activation at each individual gene locus at a given time (Figure 4F). However, in contrast with the methylation-only model, changing the methylation level by varying rates of methylation or demethylation resulted in a much more graded change in gene activation timing (Figure 4C; sensitivity coefficient = ∼0.6), even though it caused marked changes in H3K27me3 levels prior to activation (Figure 4G, H), in concordance with experimental observations (Figure 3D). This tunability was robust over different parameter ranges (Figure 4C, top right), different degrees of assembly disruption after DNA replication (Figure S2), and different degrees of cooperativity for H3K27me3 methylation (Figure S3). From a transition state theory analysis (see Mathematical Appendix and Figure S10B-C), we found that tunability arises only when interaction affinities between methylated and unmethylated nucleosomes are comparable, such that changes in methylation rates result in small changes in the height of the energy barrier to activation. Consistent with this idea, H3K27 demethylated nucleosomes still undergo PRC1-mediated compaction, and also aggregate through a variety of other PRC1-independent mechanisms (Larson et al., 2017; Strom et al., 2017). Together, these modeling results clarify how polycomb modifications can give rise to slow, tunable activation delays. First, methylation dynamics need to couple to a separate cooperative process, such as chromatin compaction. Second, compaction itself must be partially independent from methylation so that even when histone marks are depleted at the gene locus, gene repression and the compacted assembly can still be maintained via other H3K27me3-independent mechanisms (Francis et al., 2004; Larson et al., 2017; Strome et al., 2014).

### Transcription factor control of epigenetic timer function

While the stochastic time constants for *Bcl11b* activation are modulated by histone-modifying enzyme activities (Figures 3-4), our earlier work has shown that they can also be tunably controlled by two transcription factors Gata3 and TCF-1, via a far distal enhancer downstream of the *Bcl11b* gene locus (Kueh et al., 2016; Ng et al., 2018). Here, we determine whether the methylation compaction model described above can also explain tunable timing control by transcription factors in addition to chromatin-modifying enzymes. To do so, we develop an extended version of this model, where a transcription factor binds to the nucleosomal lattice, and destabilizes either the methylation or the compaction states of *N* nucleosomes within its local vicinity. Such destabilization could involve a number of candidate mechanisms: for instance, disruption of methylation could occur by direct recruitment of Kdm6a/b demethylases (Estarás et al., 2012; Seenundun et al., 2010; Williams et al., 2014), or through PRC2 eviction by chromatin-remodeling enzymes (Kadoch et al., 2017); on the other hand, disruption of compaction could involve activation of gene or lncRNA transcription (Rinn et al., 2007; Tu et al., 2017), or recruitment of factors that disrupt interactions between nucleosomes (Kraushaar et al., 2013; Talbert and Henikoff, 2017; Zhou et al., 2016).

From simulations, we found that localized disruption of a small number of nucleosomes by transcription factor binding was sufficient to enhance the rate of locus activation. Upon inhibiting compaction at a single nucleosome, there was a ∼2-fold reduction in the average waiting time to H3K27me3 loss and gene activation (Figure S4A). This time decreased further upon inhibition of compaction at additional nucleosomes. Disrupting H3K27me3 methylation at nucleosomes had a more graded effect, with perturbation of ∼10-15 nucleosomes needed for a ∼2-fold reduction in the timing of H3K27me3 loss (Figure S4B). We note that this magnitude perturbation is consistent with observed length scales for histone modification clearance around the vicinity of transcription factor binding, as measured by next generation sequencing (Hass et al., 2015; Heinz et al., 2010). Importantly, incorporation of transcription factors into this model did not affect its key dynamic properties, including its stochastic, all-or-none nature, and its ability to operate over timescales spanning multiple cell generations. Taken together, these results show that the methylation compaction model accounts for the tunability of the *Bcl11b* epigenetic timer by both transcription factors and epigenetic-modifying enzymes.

### Division independence in epigenetic timing control

How can we distinguish between the methylation compaction model, as developed above, and passive models, where H3K27me3 marks dilute out passively as a result of DNA replication? As methylation and compaction dynamics are rapid compared to cell division in the methylation-compaction model, epigenetic states at the *Bcl11b* locus recover rapidly after DNA replication (Figure 4E, S2B-C); this fast recovery, together with the invariance of chromosomal domain size with respect to cell division, could render activation timing independent of cell division speed. In contrast, in a passive dilution model, H3K27me3 loss would require DNA replication; thus, activation timing would be expected to decrease with faster cell division. To test these predicted behaviors in our models, we varied cell cycle speed in the methylation-compaction model, and measured resultant effects on activation timing. We found that gene activation timing remains largely constant, at ∼80 hr, for cell cycle durations ranging from 10-30 hours (Figure 5A right; Figure 5B). This independence between activation timing and cell cycle duration held when we adjusted the model such that DNA replication led to partial disruption of the compacted nucleosomal assembly (Figure S2). In contrast, when H3K27me3 is removed due to passive dilution by DNA replication, activation timing increases with longer cell cycle duration (Figure 5A left, Figure 5B), thus implementing a cell cycle counting mechanism for control of activation. This dependency between activation timing and cell cycle speed remained when H3K27me3 methylation and demethylation rates were non-zero, but slower or comparable to rates of cell division, such that DNA replication remained a significant factor in controlling *Bcl11b* activation (Figure S5). These simulation results provide a testable prediction to distinguish between the methylation-compaction model and passive dilution-based models for epigenetic timing control.

**Figure 5.**
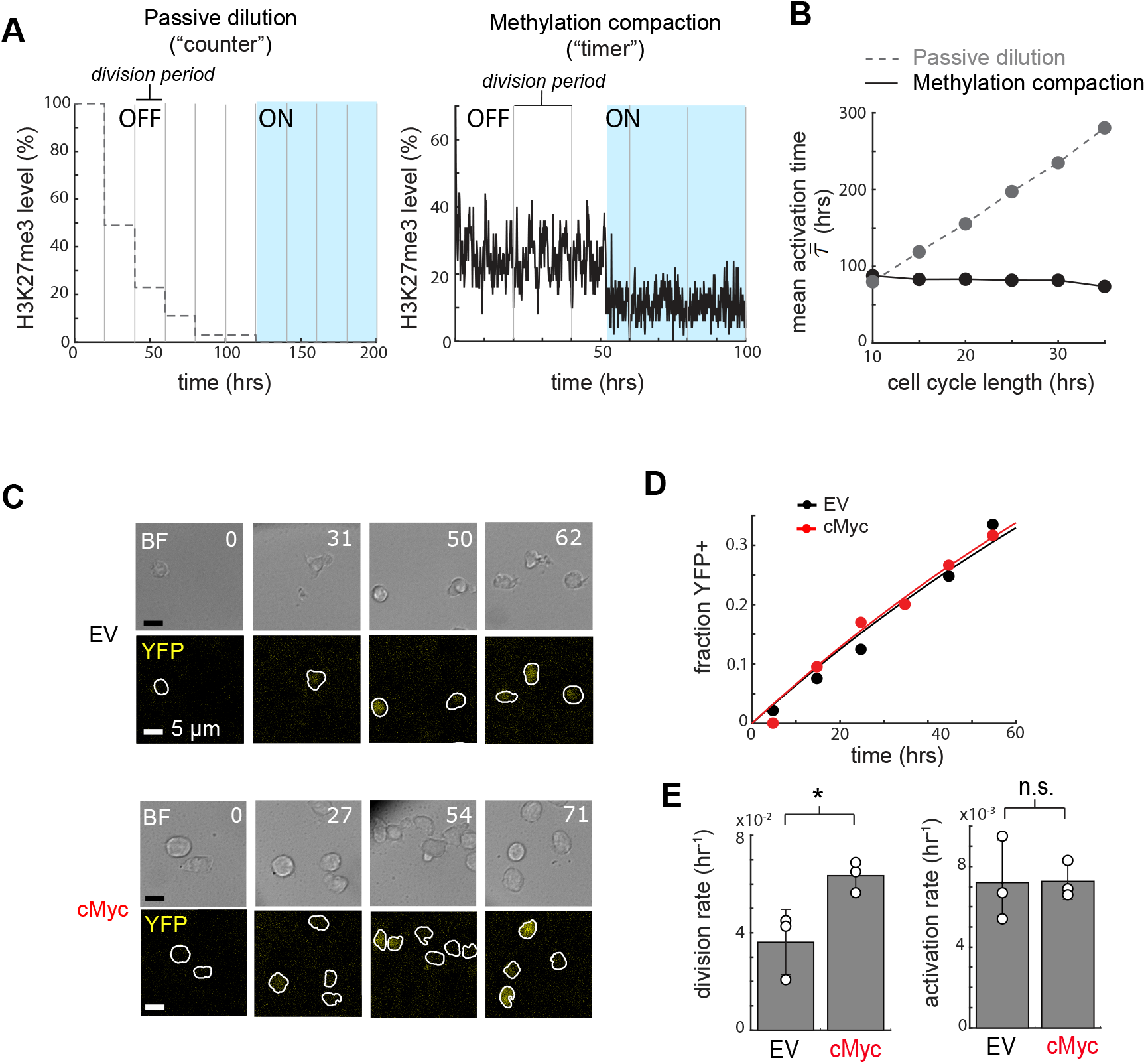
The *Bcl11b* epigenetic timer operates independently of cell-division, as predicted from the methylation-compaction model. (A) H3K27me3 time traces for a passive model, where H3K27me3 marks are diluted with each cell division (left), and the methylation-compaction model for gene activation (right). In the passive model, gene activation occurs after the total levels of repressive marks fall below a threshold. (B) Simulated average switching time as a function of cell cycle length. (C-E) Bcl11b^YFP-/RFP+^ DN2 progenitors transduced with either empty vector (EV) or cMyc CFP-expressing retrovirus were sorted, cultured on DL1-coated plates, and subject to continuous timelapse imaging. (C) Timelapse images show brightfield (BF, gray) and YFP (yellow) fluorescence of progenitors. White boundaries show automated cell segmentation. Numbers show elapsed time in hours, and scale bar = 5 μm. (D) Fraction of YFP+ cells over time. (E) Mean and standard deviation of cell division and switching rates (*n* = 3 independent experiments, *p < 0.025, n.s. not significant, one-tailed t test). Each data point is measured from one independent imaging experiment.

To test if the epigenetic timer controlling *Bcl11b* activation also operates independently of cell division, as predicted for the methylation compaction model, we altered cell division speed in DN2 progenitors by transducing them with the proto-oncogene cMyc, and then used quantitative live-cell imaging to measure resultant effects on *Bcl11b* activation. As expected, cMyc transduction increased the rate of cell division by almost two-fold compared to empty vector controls (from 0.0361 hr^-1^ to 0.0635 hr^-1^, see Figure 5C, E, left, Figure S7, Table 1 and Quantitative and Statistical Analysis). However, despite accelerating cell division, cMyc transduction did not change *Bcl11b* activation timing, with control and accelerated progenitors activating the silent *Bcl11b* allele with indistinguishable dynamics (Figure 5D), and the same time constant, as obtained by single exponential fitting (∼136 hrs) (Figure 5E, right). Taken together, these results show that *Bcl11b* activation is unaffected cell division, consistent with a methylation-compaction mechanism for epigenetic timing control.

**Table 1.** Tabulated doubling (K) and death (k_d_) rates calculated from data fitting of live and dead populations from three independent imaging experiments. Data was fitted to population dynamics model described in Statistical and Quantitative Analysis section.

|  | cMyc |  |  |  |  |  |
| --- | --- | --- | --- | --- | --- | --- |
| | Live Rate ($K$) | 95% CI | Death Rate ($k_d$) | 95% CI | Division Rate ($k_b = K + K_d$) | 95% CI |
| Trial 1 | 0.031 hrs<sup>-1</sup> | ± 0.001 | 0.034 hrs<sup>-1</sup> | ± 0.001 | 0.065 hrs<sup>-1</sup> | ± 0.001 |
| Trial 2 | 0.043 hrs<sup>-1</sup> | ± 0.001 | 0.013 hrs<sup>-1</sup> | ± 0.000 | 0.057 hrs<sup>-1</sup> | ± 0.001 |
| Trial 3 | 0.035 hrs<sup>-1</sup> | ± 0.001 | 0.027 hrs<sup>-1</sup> | ± 0.001 | 0.069 hrs<sup>-1</sup> | ± 0.001 |

|  | EV |  |  |  |  |  |
| --- | --- | --- | --- | --- | --- | --- |
| | Live Rate ($K$) | 95% CI | Death Rate ($k_d$) | 95% CI | Division Rate ($k_b = K + K_d$) | 95% CI |
| Trial 1 | 0.017 hrs<sup>-1</sup> | ± 0.001 | 0.028 hrs<sup>-1</sup> | ± 0.001 | 0.045 hrs<sup>-1</sup> | ± 0.001 |
| Trial 2 | 0.014 hrs<sup>-1</sup> | ± 0.001 | 0.007 hrs<sup>-1</sup> | ± 0.001 | 0.027 hrs<sup>-1</sup> | ± 0.001 |
| Trial 3 | 0.024 hrs<sup>-1</sup> | ± 0.001 | 0.019 hrs<sup>-1</sup> | ± 0.001 | 0.043 hrs<sup>-1</sup> | ± 0.001 |

### Temporal scalability in networks of epigenetic timers

We have described an epigenetic timing mechanism, involving all-or-none loss of a repressive H3K27me3 domain at a regulatory gene locus, that robustly generates delays in gene activation that can span multiple generations. These timing delays can be tunably controlled by both transcription factors and epigenetic-modifying enzymes. As H3K27me3 domains broadly cover lineage-specific gene loci in stem cells, the same epigenetic timing mechanism we have described could generate activation delays at other nodes in developmental gene networks; furthermore, because the polycomb repressive complexes and H3K27 demethylases have broad genome specificity, they could concurrently alter activation timing at multiple network nodes; thus, changes in their activity could lead to coordinated, scaled changes in the temporal schedules set by these networks, as is observed across different mammalian species (Figure 1).

To test this concept that epigenetic timers, when incorporated into regulatory gene networks, enable scalable control of network dynamics and developmental schedules, we modeled a hypothetical network, where an inductive signal activates a linear cascade of regulatory genes, each of which specifies a distinct differentiated cell type (Figure 6A, B, F; see Mathematical Appendix). Each gene in this cascade is activated by an epigenetic timer regulated by both an upstream regulator, and an epigenetic factor acting concurrently on all genes (Figure 6B). This gene network operates in single dividing progenitors, and as each regulatory gene in this cascade turns on, it causes the progenitor to generate the differentiated cell type specified by the activated regulator, as well as an additional copy of itself (Figure 6A). The generation of differentiated cells continues until activation of the last regulatory gene, which causes cells to terminate division and cell expansion. This asymmetric division scheme, together with the linear regulatory cascade for cell type selection, approximates the differentiation strategy in a number of neuronal systems (Kohwi and Doe, 2013; Rossi et al., 2017). As a comparison, we also analyzed a regulatory network with nodes controlled by classical gene regulation functions (Ackers et al., 1982; Bintu et al., 2005; Estrada et al., 2016), where upstream regulators activate a downstream target when their levels reach a certain threshold (Figure 6F, see Mathematical Appendix).

**Figure 6.**
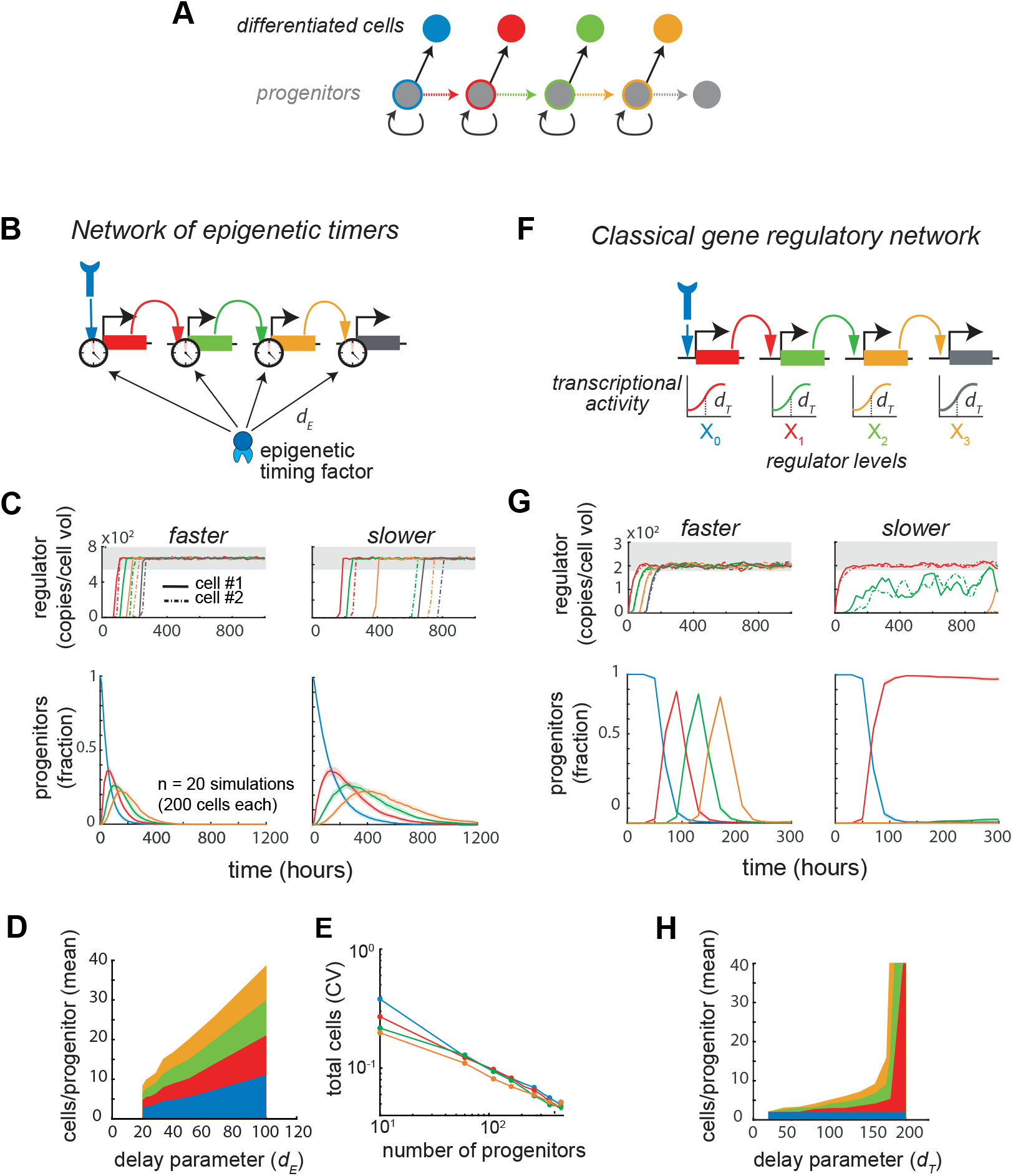
Developmental gene networks composed of epigenetic timers generate scalable temporal schedules for flexible population size control. Two gene regulatory network cascades were compared using mathematical modeling, one built from epigenetic timers (B-E), and another built from classical gene regulation functions (F-H) (See Mathematical Appendix for model description). Both regulatory networks control a common differentiation scheme (A), where progenitors that have activated a given lineage-specifying gene generate a corresponding differentiated progeny through asymmetric cell division (Rossi et al., 2017). (B) In the epigenetic timer network, where every lineage-specifying gene in the cascade is regulated by an epigenetic timing factor, whose activity is set by a common delay parameter d_E_. In the classical network (F), there is a common threshold for transcription factor activation, set by the delay parameter d_T_. (C,G, top) Stochastic simulations of the respective networks for two individual cells (solid and dashed lines) with faster (d_E_ = 25 or d_T_ = 150) and slower (d_E_ = 66.66 or d_T_ = 200) delays. Gray shaded area indicates the threshold regulator concentration required for differentiated cell production (C,G, bottom). Mean fraction of cells at each stage of the gene network with faster and slower delays. Shaded regions indicate standard deviation from 20 simulations of 200 progenitors. Fraction of cells that have turned on the last gene in the cascade is not shown. (D and H) Mean numbers of different cell types generated in the two networks, plotted as a function of the delay parameter. Proportional scaling of output cell numbers arises in the epigenetic timing network (D), but not in the classical network (H). (E) Coefficient of variation for total numbers of differentiated cells produced, plotted as a function of initial progenitor number.

By performing stochastic simulations of epigenetic timer network in single, dividing progenitors within cohorts, we found that this network could generate lineage specification schedules that could be scalably tuned by epigenetic-modifying enzymes. For a chosen set of parameters, individual progenitors turned on regulatory genes in succession upon signal exposure, with cell-to-cell variability in the timing of activation (Figure 6C, top), as expected from the stochastic nature of the epigenetic timer. Despite the cell-to-cell variability, the different activated progenitor populations within an entire cohort arose over time in a highly reproducible manner (Figure 6C, bottom), giving rise to a defined schedule for lineage specification, as well as defined numbers and proportions of differentiated cells (Figure 6D). As expected, the coefficient of variation in the numbers of different cells produced per cohort decreased steadily with increasing cohort size, indicating that this system can give rise to precise output population sizes with sufficient initial progenitor numbers (Figure 6E). Notably, this differentiation schedule was unaffected by changes in cell division rate (Figure S9A-C), indicating that the developmental network as a whole is able to operate independently from cell division. When we increased the strength of epigenetic repression by increasing the delay parameter (*d_E_*), the order of gene activation and the time duration between activation events remained the same; however, the entire differentiation schedule dilated in time (Figure 6C, bottom), allowing these progenitors to generate markedly increased numbers of different output cells, while maintaining relative proportions constant (Figure 6D).

In contrast, a network composed of classical gene regulatory nodes could not generate temporally scalable schedules. For a chosen set of parameters, this network could generate an ordered sequence for gene activation and lineage specification (Figure 6G, *slower vs faster*, and Figure S9D-F); however, when we attempted to vary activation times and population sizes by adjusting a common threshold for gene activation (*d_T_*), the dynamics of the network first showed little change, then diverged abruptly, failing to undergo the full activation sequence above a critical value (Figure 6H) This failure to fully activate caused progenitors to stall at an intermediate state, leading to disproportionate generation of early cell types and an absence of later cell types. Consistent with these observations, prior theoretical analysis had shown that count-up timers built from classical networks do not easily maintain tunable delays over multiple generations, but instead trigger activation events in a ‘now-or-never’ manner (Levine and Elowitz, 2014). Together, these results indicate that regulatory networks composed of epigenetic timers, but not those composed of classical gene regulatory nodes, can generate temporal schedules for differentiation that are robust, yet tunable over extended timescales.

## DISCUSSION

To generate organs and body plans with precise sizes and proportions, developing progenitors must follow defined temporal schedules for cell-type specification. In a range of developmental systems, these temporal schedules are upheld by cell-autonomous timers that can set and flexibly adjust developmental speeds over multiple cell generations. Here, we elucidate the mechanism of an epigenetic timer controlling the activation of *Bcl11b*, a master regulator of T-cell lineage commitment. This epigenetic timer involves the delayed, all-or-none removal of repressive H3K27me3 modifications from the *Bcl11b* locus, a switching event whose timing can be tunably controlled by the activities of opposing H3K27 methylation and demethylation enzymes. By analyzing candidate biophysical models, we find that tunable timing control arises when H3K27 methylation dynamics does not directly mediate switching, but modulates a separate physical cooperative process that can be explained by chromatin compaction. In contrast to prior models of polycomb-mediated timing control (Coleman and Struhl, 2017; Sun et al., 2014), activation delays set by this mechanism are unaffected by changes in cell cycle duration, a prediction we verify experimentally. Finally, using mathematical modeling, we find that this epigenetic timer, when incorporated into regulatory nodes of a developmental gene network, enables progenitors to adjust the speed at which they proceed through temporal schedules for lineage specification, and thereby enable them to flexibly change their output progeny numbers, while maintaining their relative proportions (Figure 6D). These results provide a solution to the long-standing question of how time is set and adjusted in developmental systems, and implicate epigenetic regulators as master controllers of developmental speed and organism size.

From analysis of candidate biophysical mechanisms, we conclude that epigenetic-modifying enzymes can tunably control activation delays only when they modulate a separate cooperative process that gates activation itself (Figure 3, 4). In standard models of epigenetic switching (Dodd et al., 2007; Zhang et al., 2014), where activation is safeguarded purely by histone methylation/demethylation dynamics, activation time constants are extremely sensitive to small changes in epigenetic-modifying enzyme activity (Figure 4A-C, left); these models cannot explain how activation times can be robustly set and tunably controlled over long timescales, as observed (Figure 3D). It is well established that H3K27me3 modifications repress gene expression by promoting chromatin compaction; furthermore, emerging evidence indicates that they do so by recruiting protein complexes that self-associate to form phase-separated condensates (Plys et al., 2018; Tatavosian et al., 2018), a physical process that can generate the cooperativity needed for all-or-none transitions in the methylation-compaction model (Figure 4E). Importantly, for activation times to be tunable in this model, nucleosomes must retain some self-interaction affinity in their demethylated states, such that methylation promotes but is not strictly necessary for nucleosomal association. This prediction is consistent with observations that nucleosomes can associate through a variety of H3K27me-independent mechanisms, involving either H3K27me3-independent binding of PRC1 to nucleosomes (Francis et al., 2004), or the self-association of other nucleosome-binding proteins (Larson et al., 2017; Strom et al., 2017). We note that this concept – that modification states of proteins modulate their interaction affinities– is well established in the study of cytoskeletal polymer dynamics (Howard, 2001; Mitchison, 1992; Phillips et al., 2012), but could provide a fresh perspective on understanding the relationship between chromatin modification states and genome structure. Further testing of the methylation-compaction model will require measurements of interaction affinities of nucleosomes in different chemical modification states, as well as identification of the molecular players that mediate these interactions. It will also require direct interrogation of the relationship between these modification states and higher order chromatin structure at individual gene loci, work that will be aided by new methods to simultaneously visualize histone modification states and chromatin folding at single gene loci in single cells (Kundu et al., 2017; Xu et al., 2018).

Our gene network modeling leads us to propose that epigenetic-modifying factors act as master controllers of developmental speed and organism size (Figure 6D-E), a prediction that is supported by evidence from a range of systems. Because of their broad sequence selectivity, epigenetic modifying factors can act concomitantly on multiple nodes in developmental gene networks (Boyer et al., 2006; Lee et al., 2006); consequently, the developmental schedules set by these networks could then be uniformly scaled by simply varying the activity of a single chromatin regulator acting globally on all network nodes. Consistent with this idea, disruptions in polycomb complex activity have been shown to accelerate differentiation across multiple cell lineages in different contexts (Akiyama et al., 2016; Endoh et al., 2017; Ezhkova et al., 2009; Fujimura et al., 2018; Jacobsen et al., 2017; Zhang et al., 2015). In particular, deletion of the PRC2 methyltransferase Ezh2 accelerates the temporal schedule for cerebral corticogenesis, leading to reduced cortical tissue size while preserving the temporal order of neuronal subtype differentiation (Pereira et al., 2010). Over evolution, such changes could have occurred through non-coding mutations affecting the expression levels of polycomb components; however, because several components of the polycomb repressive complexes, PRC1 and PRC2, have undergone extensive expansion during vertebrate evolution (Sowpati et al., 2015), it is also possible that mutations altering their protein function could have occurred to enable evolutionary timing and size variation. While H3K27me3 modifications modulate activation timing in our system, the methylation compaction model predicts that it must act with other epigenetic regulators for tunable timing control (Figure 4). This model prediction leads us to suggest that multiple epigenetic systems ultimately work together to dictate the pace of development. The repressive mark H3K9me3 also regulates lineage specification (Nicetto et al., 2019), and may co-exist with H3K27me3 to establish repression at developmental genes (Boros et al., 2014; de la Cruz et al., 2007; Yamamoto et al., 2004). Another intriguing candidate epigenetic regulatory factor is HMGA2, a chromatin-binding protein with established roles in control of body size and growth in vertebrates (Chung et al., 2018; Lamichhaney et al., 2016). HMGA proteins bind broadly to the genome to modulate chromatin architecture (Ozturk et al., 2014), and could operate broadly on many loci to control developmental speed.

The ability of this epigenetic timer to operate independently from cell division may represent a general property of developmental timing mechanisms (Burton et al., 1999; Gao et al., 1997; Heinzel et al., 2017), and may enable versatile control of organism size and form during evolution. It has long been recognized that cell proliferation rates and developmental timing can vary independently between related species to generate morphological diversity (Alberch et al., 1979). Size and shape changes can occur when timing of individual developmental events vary amid a constant backdrop of cell proliferation; indeed, there are clear examples where mutations affecting *cis*-regulatory elements alter regulatory gene activation timing to drive morphological variation (Frankel et al., 2011; Gérard et al., 1997). Conversely, morphological changes can also occur when proliferation rates vary amid a fixed temporal schedule for development (Trumpp et al., 2001). To understand how evolutionary changes generate morphological innovation in different contexts, it will be important in future studies to separately consider the impact of these changes on timing and growth during development. Hematopoiesis has long served as a convenient paradigm for studying this process. As the total number of hematopoietic stem cells appears to be roughly constant among mammals with vastly different sizes and blood production requirements (Gordon, 2002), progenitors must expand to different degrees during differentiation to generate species-specific blood cell numbers. This may be accomplished by evolutionary changes in cell-autonomous timing mechanisms that allow for more or less progenitor cell expansion prior to stochastic differentiation events. Consistent with this idea, *Bcl11b* activation and T-cell lineage commitment takes roughly twice as long in human hematopoietic progenitors when compared to those of mice under identical *in vitro* culturing conditions (Dai and Wang, 2009; La Motte-Mohs, 2005).

The need for cell-autonomous timing control is perhaps greatest in developmental systems having cells that are either spatially distributed, or interact weakly in space; thus, it is perhaps not surprising that the clearest examples for such control have thus far been found in brain and blood development. However, it is likely that autonomous timing control is also central in systems where spatial patterning controls lineage specification. While patterning mechanisms can scale cell lineage proportions according to the dimensions of a spatial domain (Ben-Zvi et al., 2011; Inomata, 2017; Rogers and Schier, 2011), they do not control the total domain size itself, which must instead be specified by different mechanisms (Averbukh et al., 2014; Fried and Iber, 2014). Moreover, from the standpoint of an organism, sizes of individual domains must be set proportionally across the entire body plan. Such proportional size scaling would be hard to achieve through coordination between different domains, but could be readily achieved through the autonomous temporal control, as we have described. For instance, cell-autonomous timers also control limb length (Saiz-Lopez et al., 2015), and may likewise be temporally scaled across different species to generate variations in length (Harrison, 1924). In general, future studies in developmental biology will benefit from closer consideration of the impact of autonomous temporal control on growth and form in multicellular organisms.

## Acknowledgements

We thank members of Kueh lab for discussions, and also thank Steve Henikoff, Michael Elowitz James Briscoe and Joe Levine for discussions, feedback, and for reading this manuscript. We also thank Steve Henikoff for the generous gift of pA-MNase for CUT&RUN experiments. This work was funded by an NIH K99/R00 Award (5R00HL119638) and a Tietze Foundation Stem Cell Scientist Award (to H.Y.K.).

## Data availability statement

The sequencing data from CUT&RUN profiling of H3K27me3 modifications will be submitted to the NCBI Gene Expression Omnibus (GEO; http://www.ncbi.nlm.nih.gov/geo/) under accession number GSE134749. The mathematical models will be deposited in the SBML BioModels Database (https://www.dev.ebi.ac.uk/biomodels/). All other data supporting the findings of this paper will be available from the authors upon reasonable request.

## Authorship Contributions

S.N., N.P. and H.Y.K. conceived the study and designed the experiments. S.N. and N.P. performed all the experiments and analyzed the data. S.N. and H.Y.K. conceived and analyzed the mathematical models. S.N. performed stochastic simulations of the mathematical models. B.I. performed analysis on imaging data. K.N. generated reporter mice for this study. S.N., N.P. and H.Y.K. wrote the manuscript.

## Methods

### Mouse Generation

*Bcl11b^RFP/YFP^* mice were generated as described before(Ng et al., 2018). Briefly, *Bcl11b^YFP/YFP^* mice were generated by inserting an IRES-H2B-mCitrine-neo cassette into the 3’ UTR of *Bcl11b* and *Bcl11b^RFP/RFP^* mice were generated by inserting an IRES-H2B-mCherrry-neo cassette into the same location. Dual allelic *Bcl11b^RFP/YFP^* mice with identical *Bcl11b* alleles except for fluorescent protein reporters were generated by breeding *Bcl11b^RFP/RFP^* mice to *Bcl11b^YFP/YFP^* mice. Bone marrow derived from F1 *Bcl11b^RFP/YFP^* mice were used for all *in vitro* T cell development assays. All Animals were bred and maintained at the University of Washington. All animal protocols were reviewed and approved by the Institute Animal Care and Use Committee at the University of Washington (Protocol No: 4397-01).

### Cell purification

To isolate hematopoietic stem and progenitor cells (HSPCs) for *in vitro* differentiation or CUT&RUN experiments, bone marrow cells were harvested from femurs and tibias of 2 to 4 month-old *Bcl11b^RFP/YFP^* mice. CD117 MicroBeads (Miltenyi Biotec) were used to enrich for HPSCs which were frozen in 90% FBS and 10% DMSO at 10^6^ cells/mL. For CUT&RUN experiments, HPSCs were further purified by staining with anti-CD117 APC-eFluor780 (ThermoFisher Scientific) and with biotinylated antibodies against a panel of bone marrow lineage markers (CD19, CD11b, CD11c, NK.1.1, Ter119, CD3ε, Gr-1 and B220 (BioLegend)). Cells were then washed with HBH (Hank Balanced Salt Solution (HBSS) with 0.1% bovine serum albumin (BSA) and 10mM HEPES) and stained with streptavidin-PerCP/Cy5.5 (BioLegend).

### *In vitro* Differentiation of T cell Progenitors

To generate double-negative (DN) T cells *in vitro*, thawed CD117^+^ cells were cultured on OP9-DL1 stromal cell monolayers as described before using standard culture medium [80% αMEM (Gibco), 20% Fetal Bovine Serum (Sigma-Aldrich), Pen-Strep-Glutamine (Gibco)], grown at 37°C in 5% CO2 conditions]. All *in vitro* T cell generation cultures were supplemented with 5ng/mL Flt3-L and 5 ng/mL IL-7 (Peprotech), and were sorted after 6 or 7 total days of culture before transducing with retroviral vectors or treating with small molecule inhibitors. DN2 cells were re-cultured in the same conditions following all cell sorting experiments.

### Flow Cytometry and Cell Sorting

Fluorescence activated cell sorting was used to isolate DN2 cells of interest with the following protocol. Bone marrow derived cell cultures were scraped and incubated in 2.4G2 Fc blocking solution and stained with anti-CD25 APC-eFluor 780 (Clone PC61.5, eBioscience) and with biotinylated antibodies against a panel of lineage markers (CD19, CD11b, CD11c, NK.1.1, Ter119, CD3ε, Gr-1 and B220 (BioLegend)). Stained cells were washed with HBH (Hank Balanced Salt Solution (HBSS) with 0.1% bovine serum albumin (BSA) and 10mM HEPES and stained with streptavidin-PerCP/Cy5.5 (BioLegend). Stained cells were washed, resuspended in HBH, and filtered through a 40-um nylon mesh for sorting with a BD FACS Aria III (BD Biosciences) with assistance from the University of Washington Pathology Flow Cytometry Core Facility. A benchtop MacsQuant VYB flow cytometer (Miltenyi Biotec) and a benchtop Attune Nxt Flow Cytometer (ThermoFisher Scientific) were used to analyze time course and perturbation experiments and acquired data were analyzed with FlowJo software (Tree Star).

### Retroviral construct and transduction

Overexpression of cMyc in DN2 cells was achieved using cMyc H2B-mCerulean MSCV retroviral vector which was described previously(Kueh et al., 2016). Retroviral mir30-based constructs (a gift from J. Zuber) were used as a backbone for delivering short hairpin RNA (shRNA)(Fellmann et al., 2013). pBAD-mTagBFP2 (a gift from V. Verkhusha, Addgene plasmid #34632) was used to substitute mTagBFP2 for the existing GFP using PCR cloning with the restriction enzymes NcoI and SalI. The RetroE-shEed retroviral construct was generated by PCR cloning as previously described(Fellmann et al., 2013) with the following PCR template: TGCTGTTGACAGTGAGCG**AAGGCATTATAAGAATAATTAA**TAGTGAAGCCACAGA TGTA**TTAATTATTCTTATAATGCCTC**TGCCTACTGCCTCGGA.

Retroviral particles were generated using the Phoenix-Eco packaging cell line as previously described(Kueh et al., 2016). Viral supernatants were collected at 2 and 3 days after transfection and immediately frozen at −80°C. To infect bone marrow derived T cell progenitors, 33 μg/mL retronectin (Clontech) and 2.67 μg/mL of DL1-extracellular domain fused to human IgG1 Fc protein (a gift from I. Bernstein) were added in a volume of 250 μL per well in 24-well tissue culture plates (Costar, Corning) and incubated overnight. Viral supernatants were added the next day into coated wells and centrifuged at 2000 rcf for 2 hours at 32°C. Bone marrow derived derived T cell progenitors used for viral transduction were cultured for 6-7 days according to conditions described above, disaggregated, filtered through a 40-μm nylon mesh, and 10^6^ cells were transferred onto each retronectin/DL1-coated virus-bound well supplemented with 5 ng/mL SCF (Peprotech), 5 ng/mL Flt3-L, and 5 ng/mL IL-7.

### CUT&RUN H3K27me3 profiling

CUT&RUN experiments were carried out as previously described^28^ with the following modifications: 1-2.5×10^5^ cells were isolated by FACS as described in sections above, bound to Concanavalin A coated magnetic beads (Bangs Laboratories), and permeabilized with 0.025% (wt/vol) digitonin. Permeabilized cells were incubated overnight at 4°C with 5ug of anti-H3K27me3 (Active Motif 39156) and then washed before incubating with protein A-MNase fusion protein (a gift from S. Henikoff) for 15 minutes at room temperature. After washing, cells were incubated in CaCl_2_ to induce MNase cleavage activity for 30 minutes at 0°C. The reaction was stopped with 2XSTOP buffer (200 mM NaCl, 20 mM EDTA, 4 mM EGTA, 50 mg/mL RNase A and 40 mg/mL glycogen) with 2pg of yeast spike-in DNA per sample. Histone-DNA complexes were isolated from insoluble nuclear chromatin by centrifugation and DNA was extracted with a NucleoSpin PCR Clean-up kit (Macherey-Nagel). For CUT&RUN quantitative PCR, human Kasumi-1 cell line (ATCC CRL-2724™) were added before binding the cells to Concanavalin A beads for internal standard instead of yeast spike-in DNA.

### CUT&RUN Library Preparation and Sequencing

Library preparation from CUT&RUN products was completed with KAPA Hyper Prep Kit (KAPA Biosystems) following standard protocol with PCR amplification settings adjusted so that annealing and extension steps are combined into one step at 60°C for 10s. Library products were size selected to be within 200 - 300 bp range using AMPure beads (Agencourt). Libraries were sequenced using an Illumina MiSeq system with paired-end 25 bp sequencing read length and TruSeq primer standard for approximately 5 millions reads per sample.

### CUT&RUN sequencing analysis

Paired-end sequencing reads were aligned separately to mouse (NCBI37/mm9) and yeast (SacCer_Apr2011/sacCer3) genomes using Bowtie2 (Langmead and Salzberg, 2012) with the following setting: --local --very-sensitive-local --no-unal --no-mixed --no-discordant −I 10 −X 700 as suggested for mapping CUT&RUN sequencing data (Skene et al., 2018). The alignment setting was designed to specifically search with high stringency for only appropriately paired reads with the proper orientation. The resulting alignments were converted to BAM files with SAMtools (Li et al., 2009) and then converted to BED files with BEDTools. Reads were sorted and filtered to remove random chromosomes. BEDTools genomecov was used to generate histograms for the mapped reads using a scaling factor that is the product of the number of spiked-in yeast reads and the number of input cells. The resulting bedGraph files were visualized using the UCSC Genome Browser (Davis et al., 2018; Kent et al.).

### CUT&RUN qPCR

Extracted DNA from CUT&RUN samples was size selected with Ampure XP magnetic beads (Beckman Coulter) to remove fragments >800bp. Primers were designed to detect the the mouse *Bcl11b* promoter (F - TCCACCTACCAGACCCCGAA, R - CTTCTTCAAAGTGCTTGGCCTC) and the human *PAX5* promoter (F - CCAGGATGTGCTGCTGTCCCAG, R - CTCCCTGGTGCTGTGCACTGA). PowerUp SYBR Green Master Mix (ThermoFisher Scientific) and CFX96 Real-Time PCR Detection System (Bio-Rad) were used for quantitative PCR. Since Kasumi-1 cells were used as internal standard, relative enrichment of H3K27me3 at *Bcl11b* was quantified by the ΔΔCq method using the human *PAX5* promoter for normalization to account for differences in efficiency and sample loss during processing.

### Cell Preparation for Time-lapse Imaging

T cell progenitors underwent *in vitro* differentiation protocol as described above. Cells were then harvested and infected with either a MSCV empty vector or cMyc overexpression vector harboring an IRES-H2B-mCerulean reporter cassette. 16-24 hours later CFP-positive cells were purified by FACS and seeded onto PDMS micromesh (250 μm hole diameter, Microsurface) mounted on top of 24-well glass bottom plate (MatTeck). To prepare the stromal-free differentiation system, the top face of PDMS micromesh was first blocked by incubating in solution of 130 μg/ml BSA while mounted on top of a 24-well plate overnight at 4°C. This step prevents subsequent binding of retronectin to the side of the meshm, allowing the cells to climb out of the microwells. Blocked micromesh was then transferred to a clean 24-well glass bottom plate. The well and mesh constructs were incubated in a solution of 10 μg/ml retronection and 3 μg/ml DL-1 overnight at 4°C. The well was then washed with PBS, and culture media [80% αMEM (Gibco), 20% Fetal Bovine Serum (Sigma-Aldrich), Pen-Strep-Glutamine (Gibco), 5 ng/ml IL-7 (Clonetech), 5 ng/ml Flt-3 (Clonetech), 50 ng/ml mSCF (Clonetech), 50 μM beta-mercapto ethanol (Sigma) grown at 37°C in 5% CO2 conditions] was added, and sorted cells were introduced at a concentration of 5-10 cells per microwell.

## Quantitative and Statistical Analysis

### Modeling simulations

All models were simulated using Gillespie algorithm provided in the Tellurium package in Python 2.7(Choi et al., 2018). Plotting of simulation results were done in MATLAB. A detailed description of the models can be found in the mathematical appendix.

### Image analysis of time-lapse movies

#### Image segmentation

Segmentation of progenitor cells were performed in MATLAB (Mathworks, Natick, MA) using custom scripts previously(Kueh et al., 2016; Ng et al., 2018). The segmentation algorithm was performed on CFP fluorescent signal as all transduced cells carried a H2B-CFP reporter cassette. Briefly, images undergo (1) correction by subtraction of uneven background signal stemming from the bottom of the glass plate or the side of the PDMS microwells (2) gaussian blur followed by pixel value saturation to fix uneven signal intensity within the nucleus of the cell and (3) Laplacian edge detection algorithm to identify the nucleus boundary. Noncell objects were excluded via size and shape limit exclusions, and segmentation parameters were chosen such that number of non-cell objects are <1% of the total segmented cells.

#### Identification of live and dead cell population

While imaging cMyc or empty vector (EV) transduced cells, we noticed that live and dead cells possess different CFP nuclear signals. Particularly, live cell nuclei have CFP fluorescence constituting a round, smooth oval shape while dead cell nuclei CFP tend to be more granular, containing distinctively small but very bright puncta. To provide unbiased recognition of live and dead cell, individual segmented cell’s CFP image patch underwent Laplacian mask filer to delineate the ‘smoothness’ of the signal and then threshold-cutoff to identify regions with high CFP signal. Resulting features such as object’s areas, perimeter, log(CFP), and puncta numbers are recoded for each cell object. Approximately 100 individual cell image patches (10% of each data set) was then manually annotated as ‘live’ or ‘dead’ by trained scientist. The results were then linked to the above feature matrix. Decision tree supervised machine learning algorithm implemented by fitctree function in MATLAB was then used to generate a model based on the annotated live/dead classification and matrix features of the training images (Fig. S6). Finally, built-in MATLAB model evaluation functions resubLoss and crossval were used to validate that mis-assignment error is below 15% for all data sets. Such approach offers an objective, automated method to distinguish between live and dead populations. All scripts for this procedure are available upon request.

### Bcl11b activation rate fitting

To measure *Bcl11b* activation rate, experimentally, Bcl11b RFP+ cells were cultured on stromal cell-free, DL1-coated system(Varnum-Finney et al., 2003), and activation of Bcl11b YFP allele was monitored in time-lapsed imaging. This stromal-free system enables a greater fold enhancement of cell division rate by cMyc transduction and better resolution for imaging as well as recapitulating Bcl11b activation and T cell lineage commitment, but supports a lower baseline rate of proliferation in unmodified cells compared to the OP9-DL1 system.

For quantitative measurement of this activation rate, first, the YFP and RFP signal intensity of segmented cells were calculated. Then each cell object underwent live/dead classification prediction by trained model as described in the previous section. Only cells that are classified as ‘live’ were selected, and their YFP RFP fluorescence 2D histogram is fitted to a two-component mixed 2D Gaussian model to obtain the fraction of YFP OFF and ON cells in the population at a given time. All the following procedures were implemented in MATLAB. Specifically:

To calculate fluorescent value of the Bcl11b YFP and Bcl11b RFP signals, pixel intensity of an annulus surrounding the segmented cell were calculated and subtracted from the raw signal intensity of the cell interior. This is to eliminate autofluorescence from the bottom of the glass plate as well as the PDMS microwells’ edge.

To obtain the time evolution of Bcl11b biallelic population fractions from initial Bcl11b YFP-RFP+ population, cells were first filtered based on their ‘live/dead’ category, and only ‘live’ cells were included in further calculation. We used a modified version of least-squares fit of a two-component mixed 2D Gaussian function described by (Ng et al., 2018) to fit the 2D histogram of Bcl11b YFP and Bcl11b RFP fluorescence levels. Specifically, let y and r be the intensity of Bcl11b YFP and Bcl11b RFP fluorescence, respectively, the overall fit, *F* (*r*, *y*), is

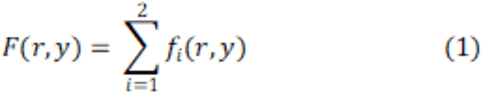

Each 2D gaussian *f* is given by:

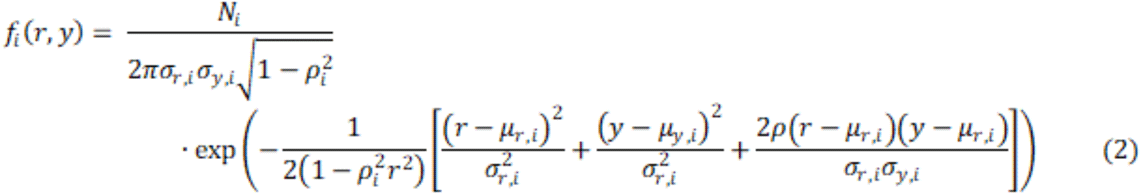

Here, *i* =1,2 correspond to the red mono-allelic and biallelic populations, since all starting cells are red mono-allelic, we excluded the other two populations (non-expressing and yellow mono-allelic). *V_i_* is the volume under the gaussian curve when integrated over r and y and is the approximation for the number of cells in each population in Bcl11b RFP mono-allelic and biallelic states.

To fit our data to *F* (*r*, *y*), we followed a two-step process described previously (Ng et al., 2018): (1) We fitted Bcl11b YFP/RFP 2D histogram at early time point (0 < t < 20) to *f*_1_(*r*, *y*) to obtain the means, standard deviations, and correlation coefficients (μ*_r_*_,1_, σ*_r_*_,1_, μ*_y_*_,1_, σ*_y_*_,1_, *p*_1_) of the Bcl11b RFP mono-allelic population. At this time point, all cells have inactive Bcl11b YFP allele. (2) Next, we fitted the 2D histograms of Bcl11b YFP/RFP levels at successive time bins of 20 hours, fixing the parameter of the first Gaussian *f*_1_(*r*, *y*), and enabling the parameters for the second Gaussian *f*_2_(*r*, *y*), to vary within bounds observed in the fluorescent distributions of Bcl11b biallelic populations. After fitting, the fraction of biallelic cell at a given time window centered on time t is given by:

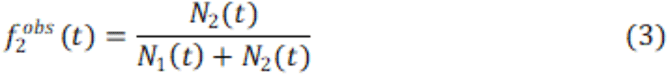

The confident bounds for *f ^obs^*(*t*) is given by:

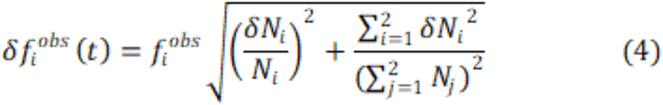

Afterward, the resulting fraction of biallelic cells as a function of time window centered at time t from the mixed Gaussian fit was then fitted to the probability density function of a first order process:

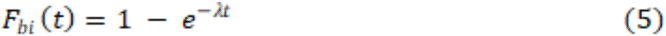

Where is the activation rate of Bcl11b from RFP-monoallelic to biallelic state. We chose this function for activation rate fitting since our histone dynamics simulations suggested Bcl11b activation can be estimated as a first order stochastic process (see Fig. 4-5). For this function, fitting was done using MATLAB fit function, and 95% confidence interval for the fit was recorded.

### Population dynamics model and fitting

We built a mathematical model to describe the population dynamics of progenitor cells transfected with empty vector (EV) and cMyc. From initial inspection of time-lapse movies (Fig. 5C), progenitors transduced with cMyc appear to expand more quickly than control progenitors, as expected. Faster expansion of cMyc-transduced cells could be due to faster cell cycling or slower cell death. To disentangle these two effects, we quantified numbers of both live and dead cells over time (Fig. S7) and fit these data to population dynamics models to obtain division and death rates:

In general, the model includes a population of live cell (*X*) with a birthrate *k_b_* and a death rate *k_d_* to generate the dead cell population (*Y*). This population in turn has a clearance rate δ denoting the process in which CFP level degrades and the dead cells become undetected.

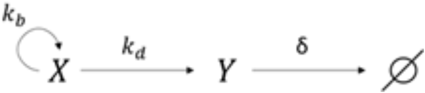

Live population at a given time *T* can be described as a simple first order process:

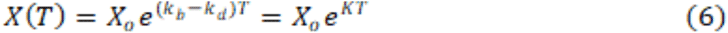

Where *K* = *k_b_ − k_d_* and *X_o_* is the initial number of live cells at the start of imaging. On the other hand, we adopt a stepwise approach to model the dead cell population:

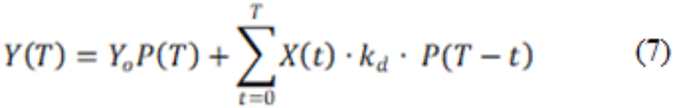

Here *Y _o_* is initial number of live cells at the start of imaging and *P* (τ) is an exponential decay function describing the fraction of CFP-positive dead population remaining after a period time from its first appearance. Since dead cell’s fluorescent signal is dim, segmentation of these cells tends to ‘fickle’ before completely disappear. In this model, whenever a cell starts to die, its probability of being detected decreases as per function *P* (*T*), and the number of dead cells at a given time T is the sum of all the still-detected dead cells generated since the start of imaging up until T. This discrete approach allows us to fit cell death data to a relatively simpler function compared to a more sophisticated two-component system of ODE model without sacrificing the complexity of dead cell detection phenomenon.

We determined *P* (τ) empirically for EV and cMyc population separately by manually following 30 different dead cells and record the time period in which it was detected and undetected until complete disappearance. Then we plotted how many dead cells out of 30 were detected after a given time has elapsed. An exponential function decay function was used to fit this ‘fraction detected’ curve and to estimate value for clearance rate (Fig. S7B):

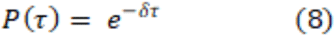

Here, *P* (τ) is the probability of a given dead cell to be detected under the CFP fluorescent channel after period of time since its initial death. δ is the clearance rate of this process.

To fit imaging data to equation (6). We classified segmented cell object as live or dead using trained machine learning model as described in ‘Image analysis of time-lapse movies*’* section. Number of live cells as a function of time was fitted to equation (6) using MATLAB fit function to estimate, and 95% confidence interval for the fit was recorded.

To fit imaging data to equation (7), we performed fine scanning of candidate *k_d_*_,*i*_ values from the set *K_D_* = {0, 0.0005, 0.001, 0.0015, …, 0.0495, 0.05}. For each *k_d_*_,*i*_, a predicted *Y _p_*_,*i*_(*T*) curve was generated based on equation (7) where *T* = *t*_1_, *t*_2_, *t*_3_, … with *t_i_* being the time point at which experimental measurement took place. *Y _p_*_,*i*_(*T*) is then compared to the experimentally observed dead cell number *Y _exp_*(*T*) using sum square error method:

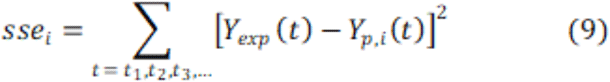

The best fit *k_d_* value is chosen to be the *k_d_*_,*i*_ value whose *sse_i_* is the smallest.

In order to calculate the confidence bound of the fit, we utilized nonlinear regression method by first calculating the residue of the model’s predicted values *Y _p_*(*t_i_*):

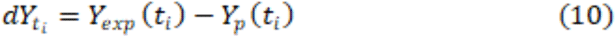

Then we calculate to the Jacobian of the model function to estimate the covariance at each time point and is given by:

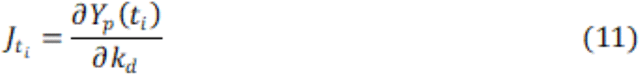

These inputs were used to estimate 95% confidence interval using MATLAB ‘Nonlinear regression parameter confidence intervals’ function nlparci.

Summary of results from fitting of data to activation rate and population models are tabulated in Table 1.

---

**Supplementary Figure 1.**
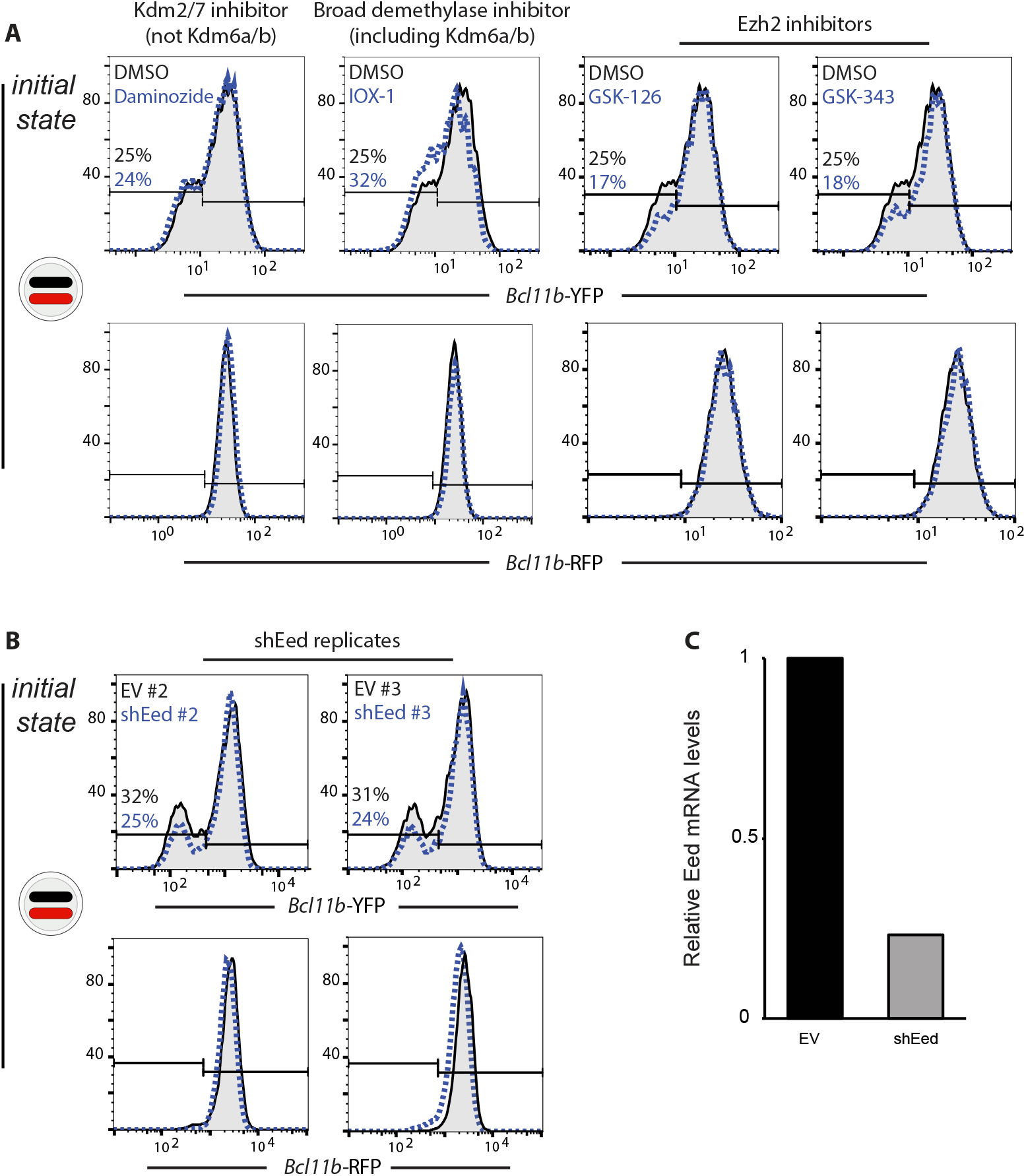
H3K27me-modifying enzymes modulate *Bcl11b* activation timing. (A) DN2 Bcl11b-RFP monoallelic cells were sorted, recultured in the presence of different small molecule inhibitors and analyzed by flow cytometry 3 days later. Slowed activation rate is unique to Kdm6a/b inhibitors: IOX-1 (above) or GSK-J4 (Figure 3D). Similar accelerated activation rate was observed for all three Ezh2 inhibitors: GSK-126, GSK-343 (above), and UNC1999 (Figure 3D). (B) Two additional experimental replicates for Eed knockdown show similar accelerated activation rate (Figure 3D). (C) Relative mRNA levels of Eed were measured by qPCR.

**Supplemental Figure 2.**
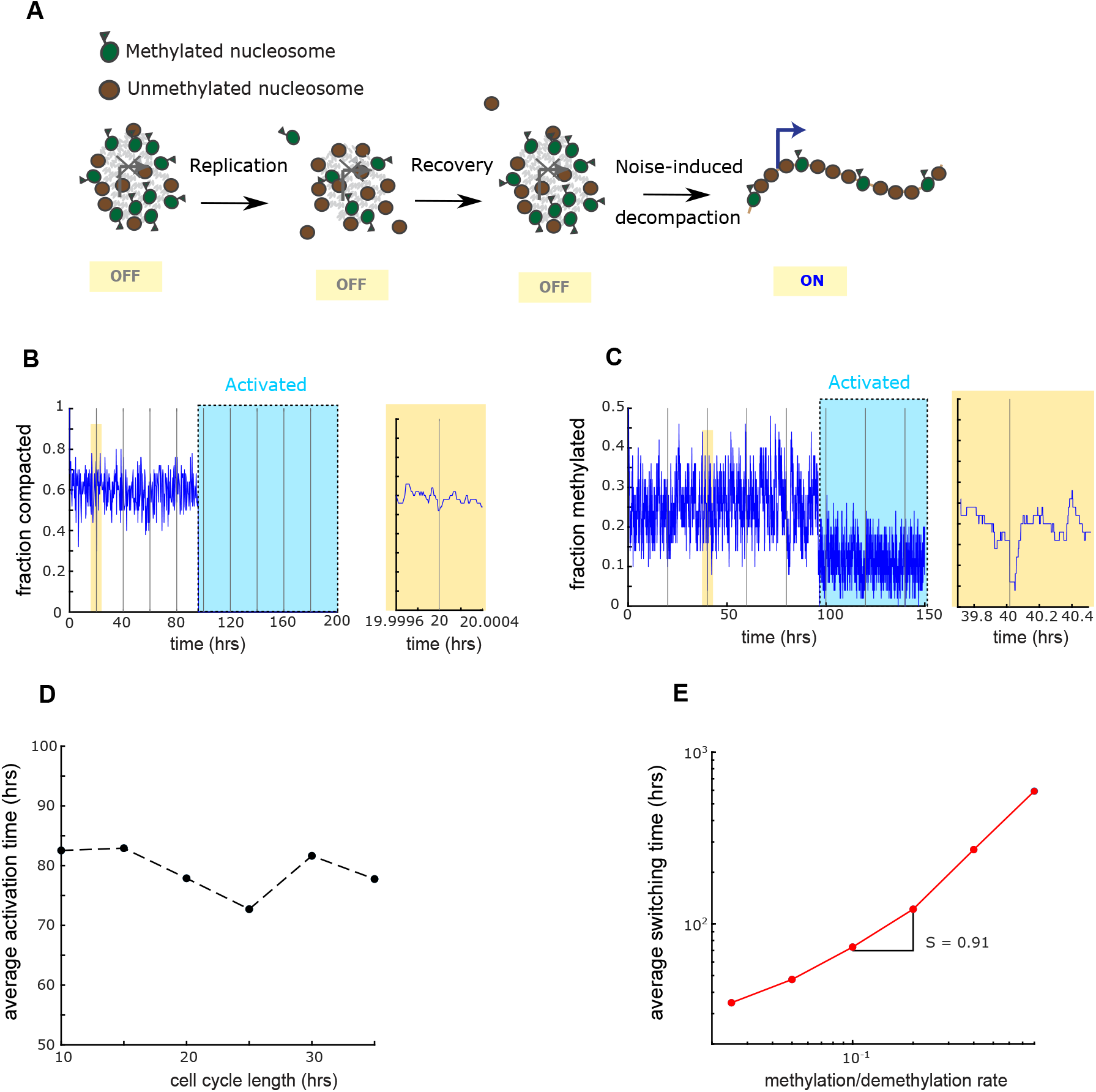
Perturbations to the compacted state by DNA replication does not affect tunability or division-independence in the methylation compaction model. (A) Modified methylation compaction model where every cell division leads to 50% reduction in methylation state and 10% reduction in compaction state. (B-C) Model’s compaction and methylation state as a function of time. Zoomed in first replication event. (D) Average switching time of the system as a function of cell cycle length. (E) Model systems’ average switching times as a function of methylation and demethylation rate ratio. Tunability coefficient S (Δ logY/ΔlogX) for each plot was calculated by taking the slope of the linear fit y = ax+b for the methylation model data set.

**Supplemental Figure 3.**
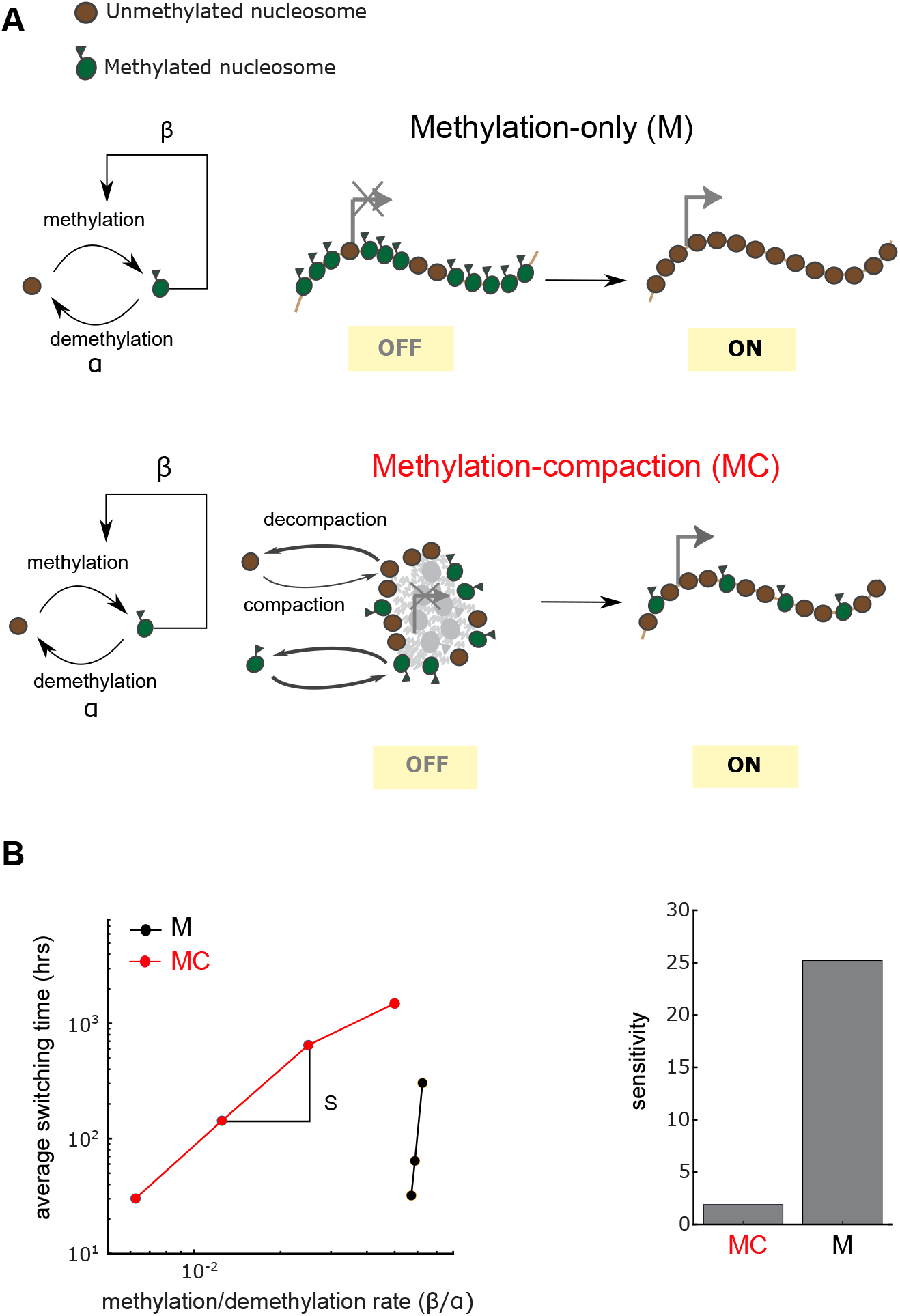
A methylation-compaction mechanism, with cooperative methylation dynamics, also shows enhanced tunability in activation timing compared to the methylation-only model. (A) Top - Methylation model enables gene activation via complete eviction of methylation marks. Bottom - Methylation compaction model with cooperative methylation rate. A nucleosome’s methylation rate increases with the number of methylated nucleosomes in the system. (B) Average switching times as a function of methylation (β) and demethylation rate constant (α) ratio for the methylation (black) and methylation compaction (red) models. Tunability coefficient (ΔlogY/ΔlogX) for each plot was calculated by taking the slope of the linear fit y = ax+b for the methylation model data set and the last 5 data points for the compaction model.

**Supplementary Figure 4.**
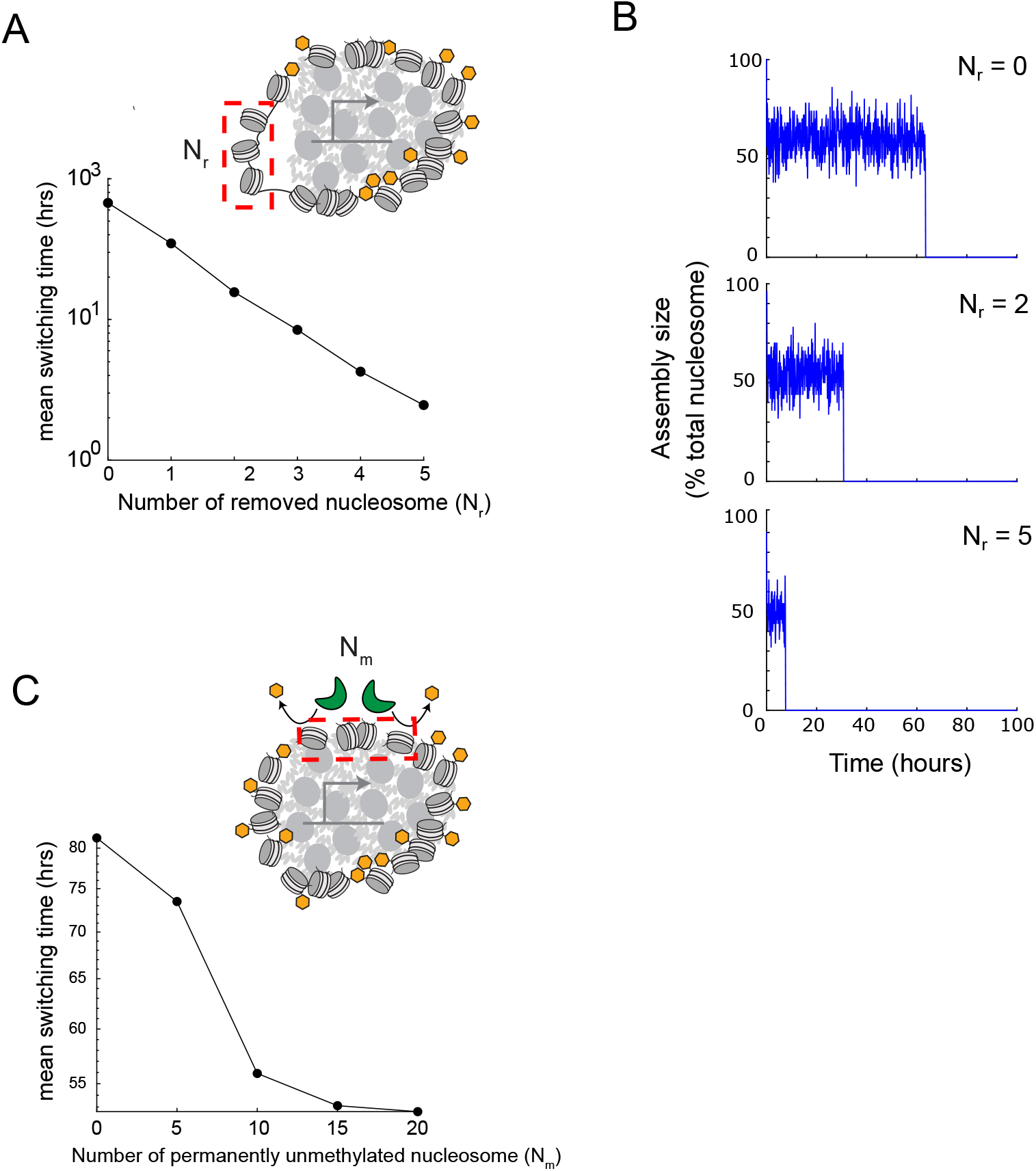
Local disruptions in nucleosomal methylation or compaction, caused by transcription factor binding, generates tunable changes in activation time. Simulations of the methylation-compacted mechanism were performed where a fixed number of nucleosomes were either blocked from entering a compacted assembly (A,B), or from acquiring H3K27me3 marks (C). (A) Mean activation timing of the methylation compaction model (MC) as a function of number of removed nucleosome from the original domain size of 50. B) Time evolution of compaction state in one sample simulation with indicated number of removed nucleosome. C) Mean activation timing of the MC model as number of non-methylatable nucleosome increases.

**Supplementary Figure 5.**
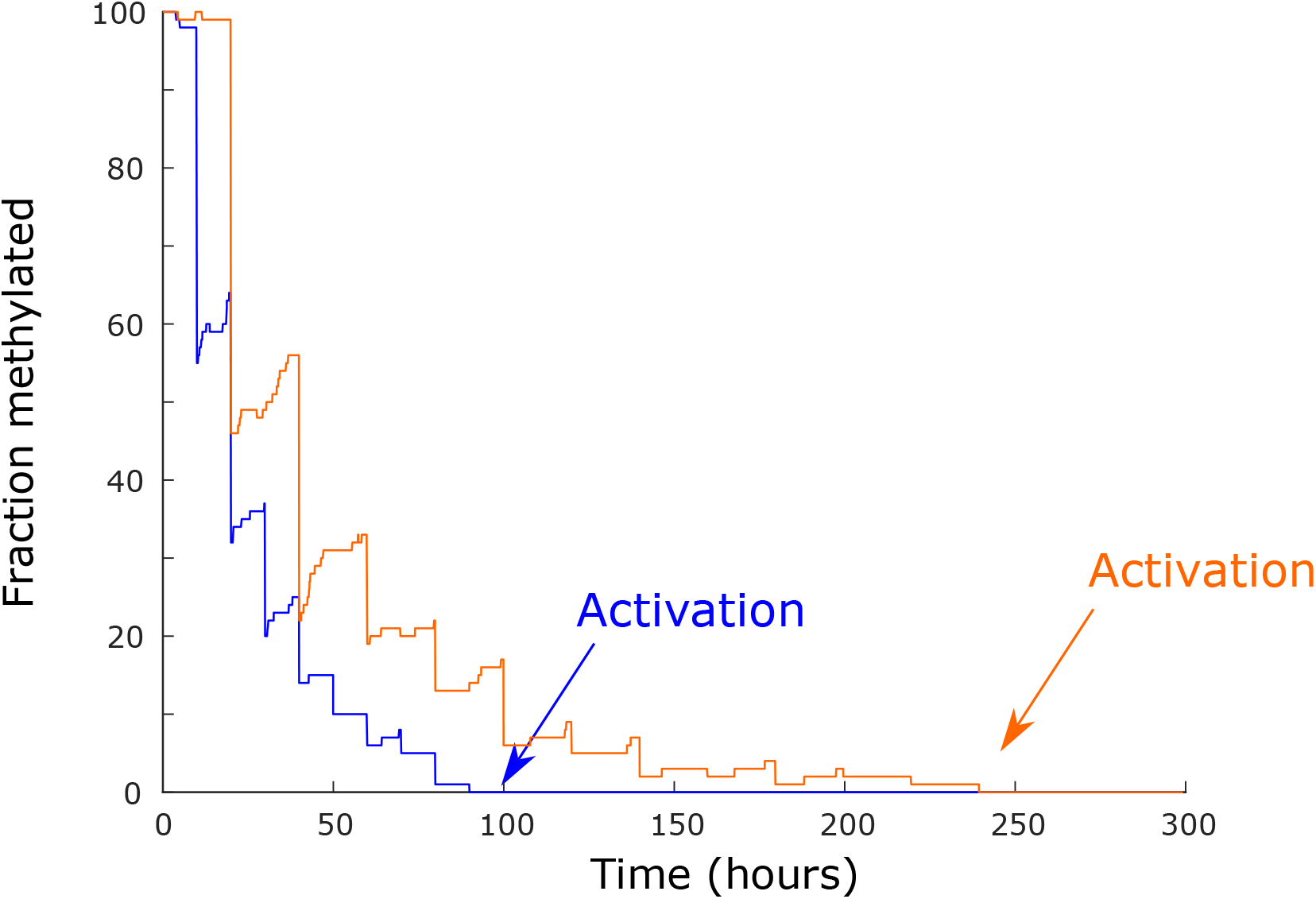
Modeling of H3K27me3 passive dilution with minimal methylation and demethylation shows a significant dependence of activation time on cell cycle length. Fractions of methylated histone are shown for the methylation only model with cell division lengths set to be 10 hrs (blue) and 20 hrs (orange). Methylation and demethylation rates were set to 0.001 per hour (See Mathematical Analysis for more detail).

**Supplementary Figure 6.**
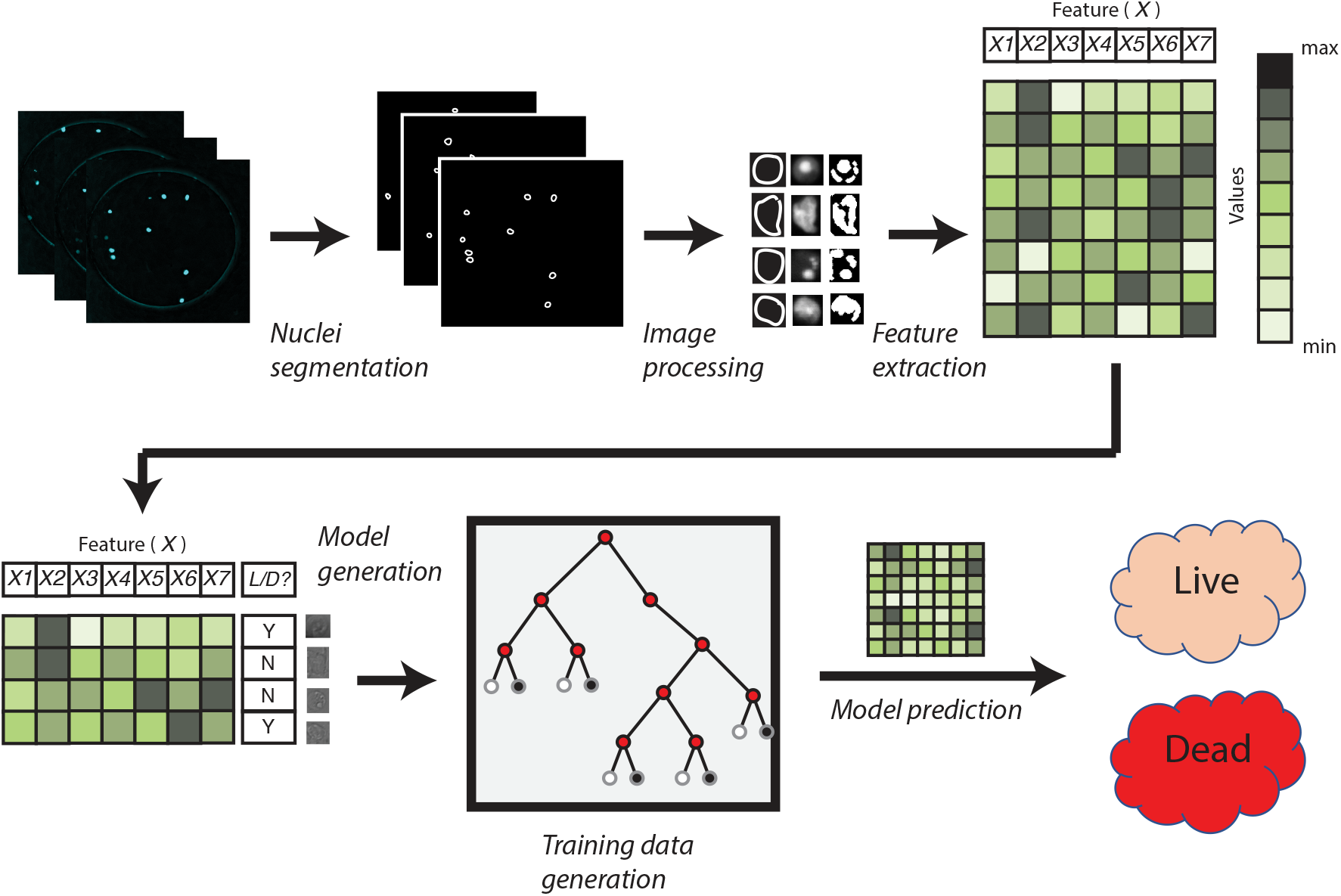
Image processing pipeline for identification of live and dead cells in imaging experiments. Cyan fluorescent protein (CFP) signals from transduced cells are used for segmentation of individual cell. Segmented populations are analyzed using imaging processing tools in MATLAB, and specific features such as object area, perimeter, and CFP intensity are collected. Approximately 10% of the total individual cell are then manually labeled as live or dead and fed into classification tree machine learning algorithm to generate a classification model. The rest of segmented cells are classified subsequently via the trained model.

**Supplemental Figure 7.**
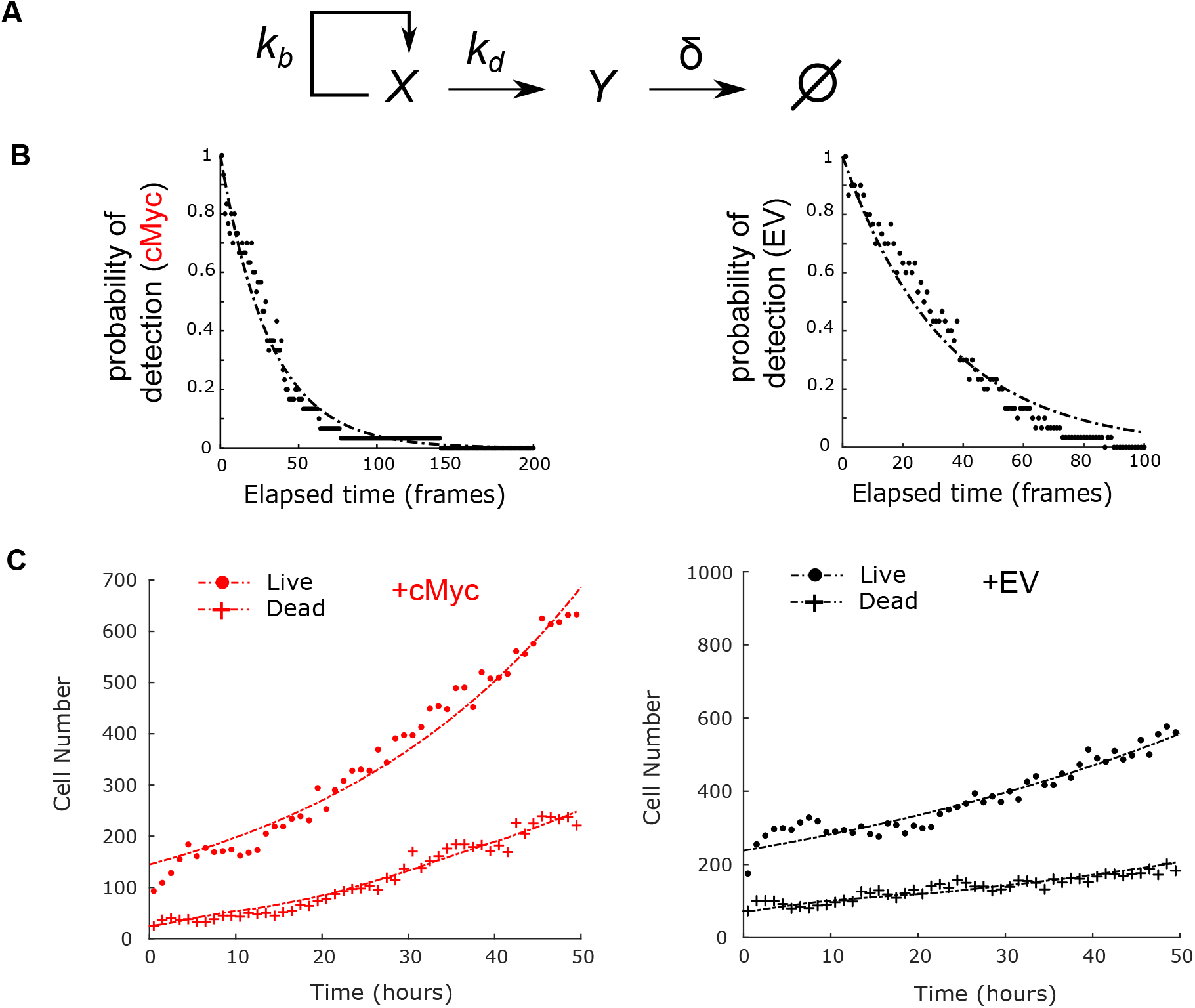
Mathematical model for disentangling cell division and death rates in cells subject to timelapse imaging. (A) Mathematical model describing the population dynamics of dual-color Bcl11b cells transfected with cMyc and empty vector (EV). The model includes a population of live cells (X) with a birthrate k_b_ and a death rate k_d_ the generate the dead cell population (Y). This population then has a permanent clearance rate of δ, indicating the process in which these cells’ CFP degrade, and the dead cells become undetected. (B) Experimentally determined decay of dead cells’ detectability in time-lapsed movies. Individual dead cells were followed until their CFP level completely diminished to the point of undetection by the segmentation algorithm, and the elapsed time was recorded. Thirty different individuals cells were recorded for each cMyc and empty vector populations. Data was fit to an exponential decay function P(τ) = e^-δτ^ with δ = 0.032 per frame for cMyc and δ = 0.030 per frame for EV population. (C) Representative time evolution of cell number in live and dead cell populations in one live-imaging experiment. Dual-color Bcl11b DN2 progenitors were transfected with cMyc and empty vector expressing retroviral vector. Transfected Bcl11b YFP-/RFP+ cells were cultured on DL1 + retronectin coated plates and activation of YFP allele was monitored. Representative live and dead population dynamics of cMyc (red) and empty vector (black) populations from one movie were shown on the right. Data was fitted to population dynamics model described in (A).

**Supplemental Figure 8.**
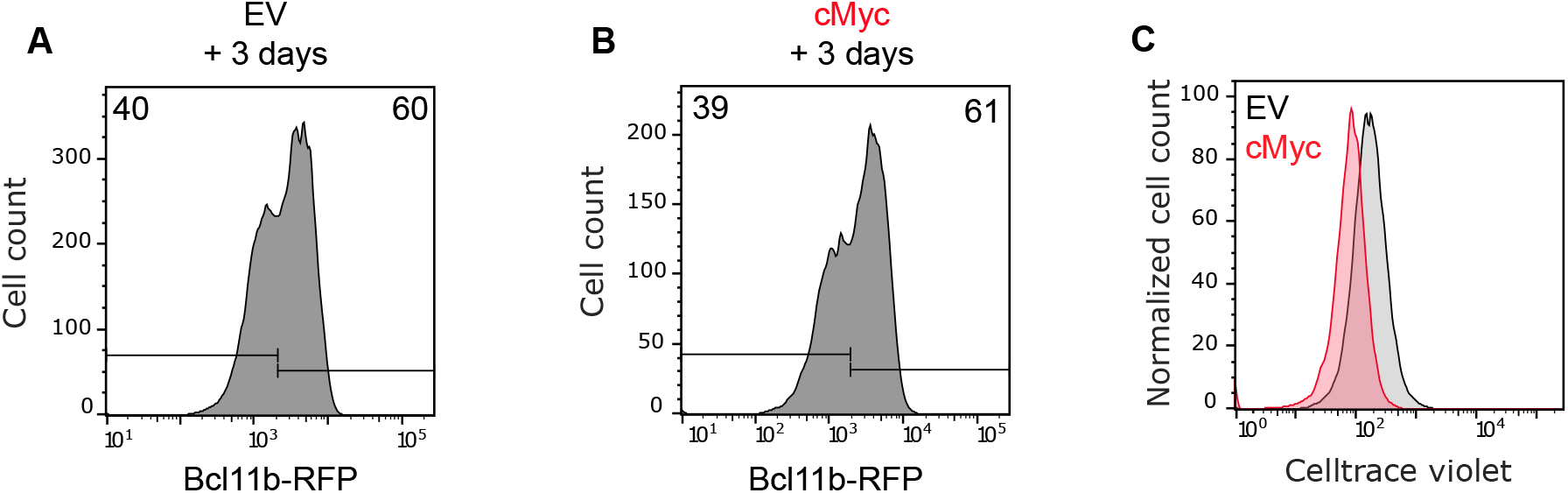
B*c*l11b activation timing is independent of cell division rate in OP9-DL1 culture system. *Bcl11b*^YFP+/RFP-^ DN2 progenitors were transduced with a CFP-expressing empty vector (A) or cMyc (B) retrovirus, sorted and cultured on OP9-DL1 stromal layers with 5 ng/mL IL-7 and Flt3L. RFP allele activation was analyzed 72 hours later via flow cytometry. Plots show similar Bcl11b activation percentages in EV versus cMyc-transduced cells. (C) CellTrace Violet profiles of cMyc (red) and empty vector (grey) population confirm faster division rate of cMyc overexpressing cells.

**Supplementary Figure 9.**
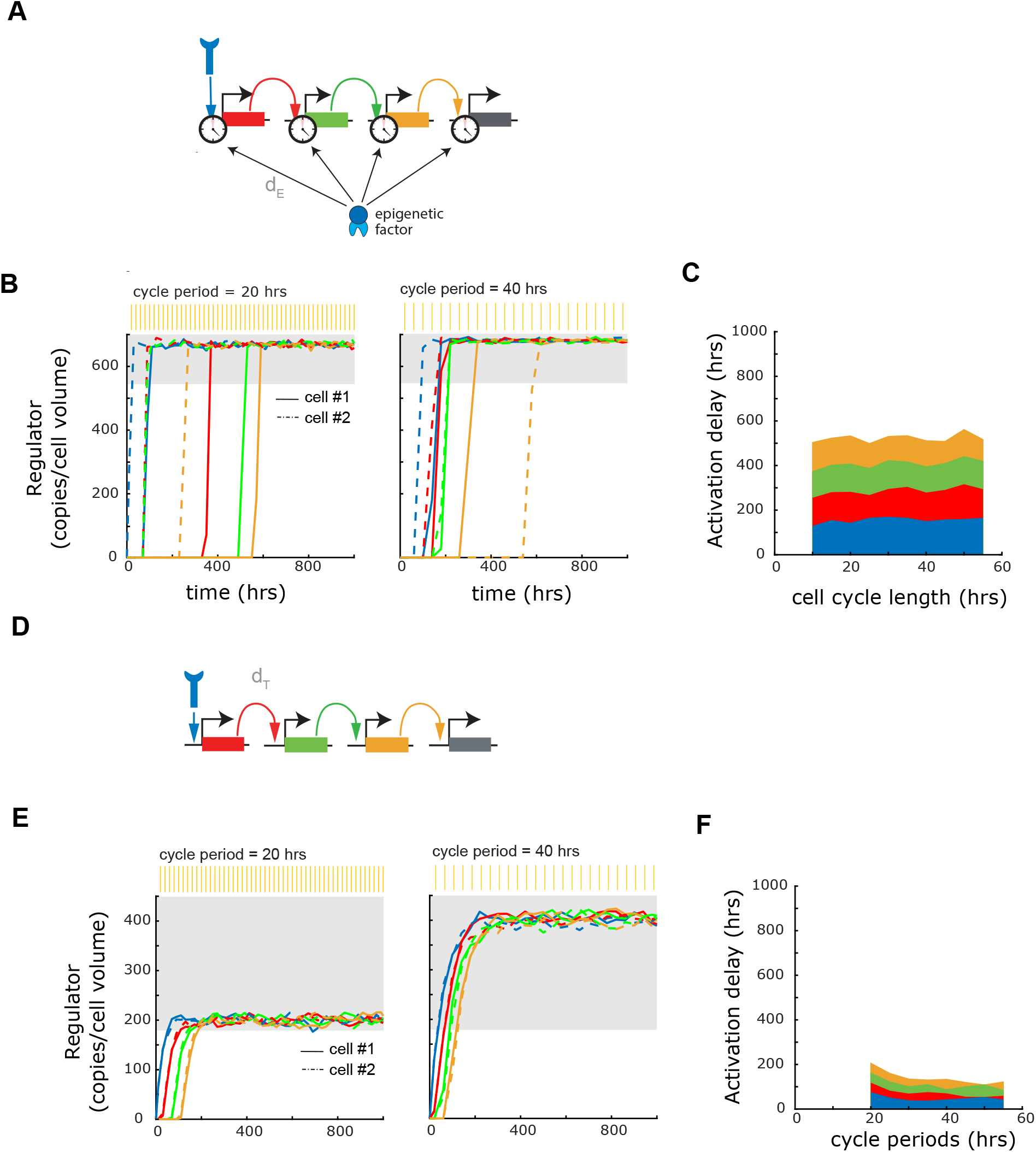
Developmental gene networks built from epigenetic timers can set temporal schedules that are independent of cell division. Cascade of activated genes each controlled by an polycomb-based timer (“E”) with a delay parameter d_E_ (A) or a classical regulation timer with a delay parameter d_T_ (D). (B and E, top) Stochastic simulations in two cells (solid and dashed lines) with faster (once every 20 hrs, left) and slower (once every 40 hrs, right) cell cycling rates. (C and F) Activation delay, Activation delay for each gene, defined as the time it takes for 85% of cells to reach its “ON” state in a given population, as a function of cell cycle period.

**Supplementary Figure 10.**
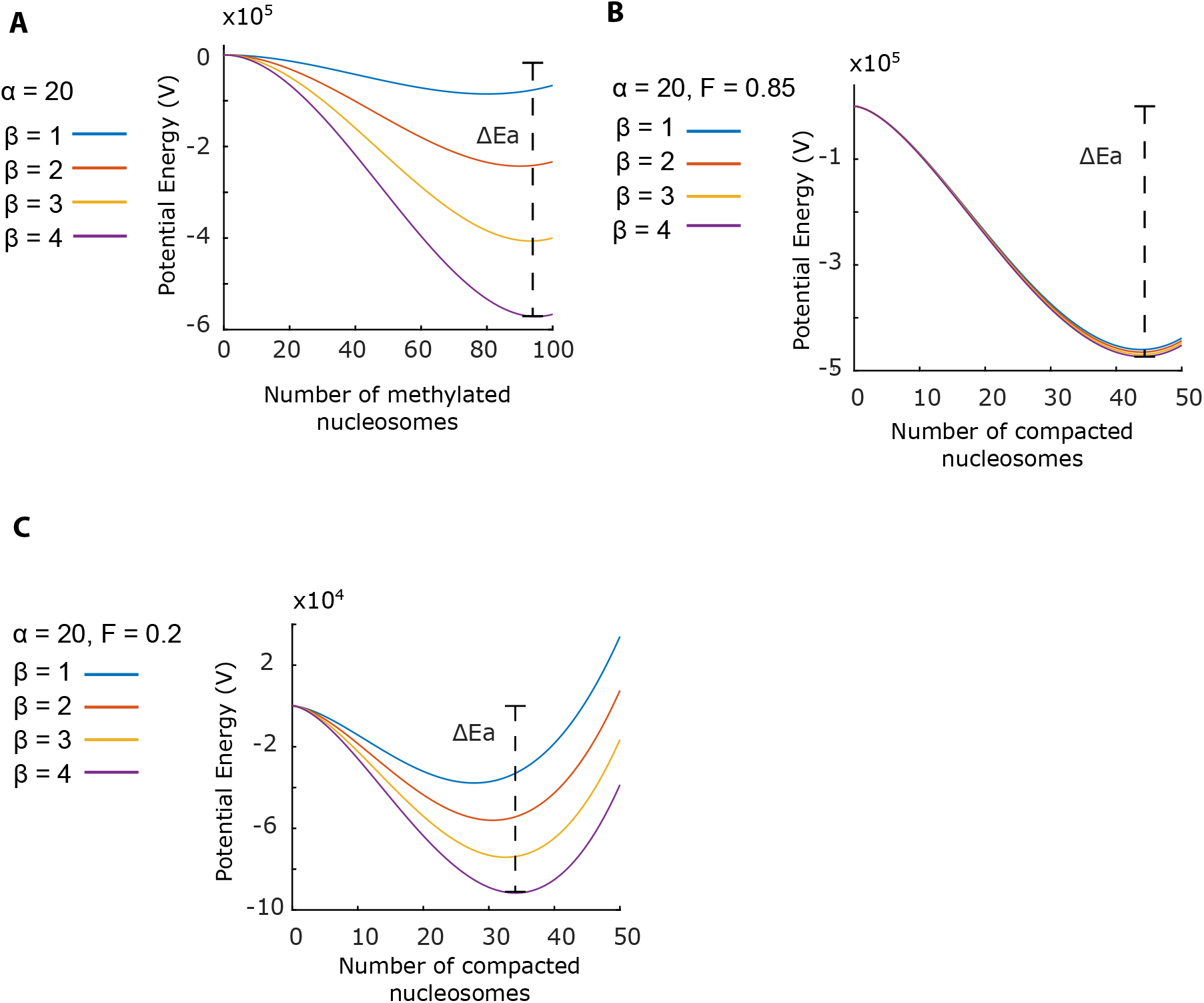
Activation energy is robust to changes in methylation rate when interaction affinities between methylated and demethylated nucleosomes are similar. (A) Potential energy landscapes of methylation model (M). (B-C) Potential energy landscapes of the methylation-compaction model (MC). Parameter Fdictating the compaction affinity reduction in demethylated nucleosome in the MC model was set to 0.85 (B) and 0.1 (C). Activation energy barrier (Ea) is defined as the potential energy (V) height between the local maximum and local minimum of the potential energy landscape. Each potential curve was plotted with demethylation parameter = 20 hrs^-1^ and methylation rate parameter as indicated by the curve’s color (See Mathematical Appendix).

## MATHEMATICAL APPENDIX

### Part I: Modeling of the polycomb-dependent epigenetic timer

#### Introduction

To understand the epigenetic timer controlling the *Bcl11b* activation, we used mathematical modeling to analyze a series of candidate biophysical mechanisms. In this modeling, we seek to explain the central emergent properties of the switch, namely (1) its irreversible, all-or-none nature; (2) its long, stochastic time delay; (3) the heritability of its inactive and active states over DNA replication; and (4) its tunability with respect to changes in H3K27me3 levels and modifying-enzyme activity.

We consider two candidate mechanisms. In the methylation only mechanism (Model I), individual nucleosomes within in a one-dimensional lattice can be methylated or unmethylated. Gene expression is assumed to occur when the total fraction of methylated nucleosomes in this lattice falls below a threshold value. In the coupled methylation compaction mechanism (Model II), individual nucleosomes are also methylated and demethylated; in addition, these nucleosomes also interact to form a compacted assembly with rates dependent on the H3K27me3 marking. Unlike Model I, gene expression does not depend directly on H3K27me3 levels, but on the compaction state of the nucleosome assembly, which in turn depends on methylation states of individual nucleosomes. Both models explicitly model DNA replication as a process involving random segregation of modified nucleosomes into daughter strands. From our analysis, we find that the methylation-compaction mechanism alone explains the observed emergent behaviors of the *Bcl11b* epigenetic timer, and thus represents our favored model for its underlying mechanism.

#### Model I: The Methylation Only Mechanism (M)

We adopt a standard framework for histone modification dynamics previously shown to generate multistability (Angel et al., 2011; Dodd et al., 2007). In this model, individual nucleosomes reside in a one-dimensional lattice, and exist in two states, a methylated state, corresponding to an H3K27 tri-methylated state, and demethylated state. We do not describe multiple demethylated states in our model (i.e. mono-methylation, di-methylation, and an un-methylated state), though our theoretical analysis, together with previous work (Dodd et al., 2007), indicates that our main conclusions hold in more complex models with additional states. As with previous models, the methylation rate of a given nucleosome depends on the number and distance of methylated nucleosomes in its vicinity, reflecting observations that PRC2 can bind and be activated by H3K27me3-marked nucleosomes to write H3K27me3 on neighboring nucleosomes. Demethylation is taken to occur at a first order rate. In this model, we assume there is no spontaneous methylation in the absence of existing methylated nucleosomes; thus, once all nucleosomes are demethylated, no more remethylation is possible and the system enters an irreversibly activated state.

##### Methylation

We explicitly model mark binding and cooperative activities of the PRC2 complex, as well as the methylation state of each individual nucleosome. Let *p_i_* be the methylation state of the *i*th nucleosome. *p_i_* = 0 denotes the de-methylated state while *p_i_* = 1 denotes the methylated state.

Let *u*′ and *u* denote the transitions between the methylation state and demethylation state, respectively. The model is set up as follows:

For i ∈ {1, . ., 100}:

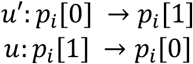

With:

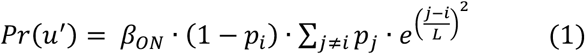

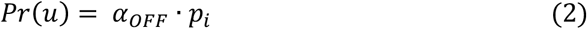

The parameter *L* can be interpreted as the ‘reach’ of the PRC2 complex to neighboring nucleosomes. Large value of *L* indicates a long length scale for interaction with nearby nucleosomes. This effect is set to have a gaussian shape so that nucleosome closest to the anchored PRC2 complex has the highest methylation rate. Similar distributions of activity have been reported for artificially tethered enzymes (Hass et al., 2015), as well as for histone modifications around transcription factor binding sites (Heinz et al., 2010). Moreover, we assume periodic boundary conditions for the one-dimensional lattice, though similar results were observed with other non-repeating boundary conditions (not shown).

##### Cell division

To model the transmission of histone marks across cell divisions, we assume that methylated nucleosomes segregate randomly to the two daughter DNA strands upon replication; thus, each nucleosome position has a probability *p =* 0.5 of inheriting a nucleosome that is methylated. Experimental evidence suggests that approximately 50% of total global H3K27me3 partitioning of parental marks to the subsequent generations (Alabert et al., 2015).

From stochastic simulations of this model, we find that this methylation only mechanism can generate a slow, heritable, and stochastic gene switch (see Results and Fig. 4); however, switching times are hypersensitive to mild changes to methylation and de-methylation rates, and therefore inconsistent with experimental results. To understand the origins of this hypersensitivity, we re-formulate this model using a chemical kinetics framework amenable to analysis using transition state theory. By considering the limit where *L* → ∞, such that each H3K27me3-bound PRC2 methylates all other un-methylated nucleosomes with the same reaction rate. In this limit, given *N* as number of unmethylated nucleosome, we can completely describe the state of the system by the number of methylated nucleosomes *N’*. As the rates of adding or subtracting one methylated nucleosome from the system would reduce to become a function of *N’*, independent of spatial arrangement:

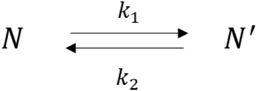

Where

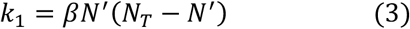

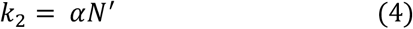

and N*_T_* is the total number of nucleosomes. The master equation describing the time evolution of this system is given by:

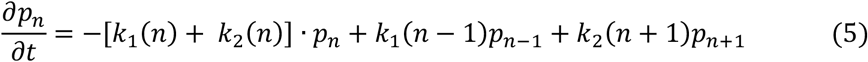

where *p*_*n*_ is the probability of having *N’* methylated nucleosomes. When the total number of nucleosomes is large, we can approximate the number of methylated nucleosomes to be a continuous variable \. In this limit, we can rewrite the master equation as Fokker-Planck equation:

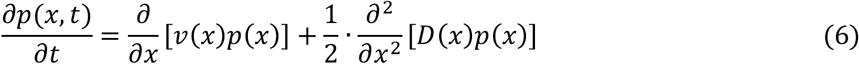

where, we have ignored third and higher order terms, and where:

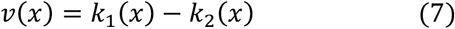

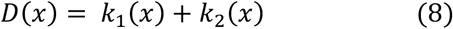

Given the velocity and diffusion constants for this system as a function of methylated nucleosome number, the switching of the system is essentially given by the first-passage time of the system to reach the absorbing state *x = 0*. However, a closed-form solution of this first-passage time distribution for the given rate functions is hard to obtain; Nevertheless, we note that our system operates in the regime where the timescales of individual methylation and demethylation reactions are much shorter than switching times for this system. In this regime, switching times are well described by the Kramer’s theory for escape of a Brownian particle over a potential well (Kramers, 1940), and would thus approximately scale exponentially with the height of a potential energy barrier. We can obtain the functional form of this potential barrier by relating it to the velocity function:

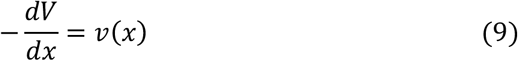

From equations (3), (4), and (7), we can then integrate the system to explicitly derive the potential function:

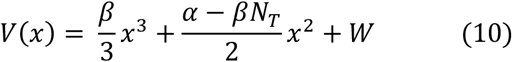

Where *W* is an arbitrary number. A plot of *V*(*x*) is demonstrated in Fig. S10A. The energy landscape possesses a local minimum at a nonzero value of *x*, indicating the metastable state. The landscape has a local maximum near *x* = 0. State switching occurs when the system reaches the absorbing state *x = 0*. Thus, we define *E*_*a*_ as the height in *V*(*x*) between the local maxima and the metastable state minima. When we plotted this potential energy for different values of β (Fig. S10A), we found that moderate changes in β led to significant changes in potential well height. As switching rates scale roughly exponentially with well height, we would expect this system would show extreme sensitivity in switching times with respect to changes in methylation rate changes.

### Model II: The Methylation Compaction Mechanism (MC)

As the methylation only mechanism above does not fully explain the tunable characteristics of the *Bcl11b* activation timer, we considered a second model, where histone methylation is coupled to chromatin compaction. Indeed, H3K27me3 has been suggested to meditate a condensed, polymerase-inaccessible chromatin conformation. For instance, the polycomb repressive complex 1 (PRC1), which binds H3K27me3 and is important for compaction and gene silencing, oligomerizes through contacts on its Bmi1 or Phc subunits (Eskeland et al., 2010; Gray et al., 2016; Isono et al., 2013; Kahn et al., 2016), and may also undergo weak, multivalent interactions on Cbx2 that result in liquid-liquid phase separation and gene silencing (Howard, 2001; Larson et al., 2017). This model consists of two main modules: (1) a H3K27 methylation and demethylation mechanism, and (2) a chromatin decompaction mechanism linked to H3K27me3 modifications, that ultimately underlies gene switching. In our description of compaction dynamics, we do not explicitly model the spatial extent of the compacted assembly; instead, we adopt a mean-field approach that is established in models of cytoskeletal polymer dynamics (Erickson and Pantaloni, 1981; Jackson and Berkowitz, 1980). With this approach, the numbers of un-methylated and methylated nucleosomes within a compacted assembly are given by C and C’ respectively, along with those outside the assembly are given by D and D’ respectively. As a result, the dynamical system is described by four states: 1) Compacted-Methylated 2) Compacted-Demethylated 3) Decompacted-Methylated and 4) Compacted-Demethylated:

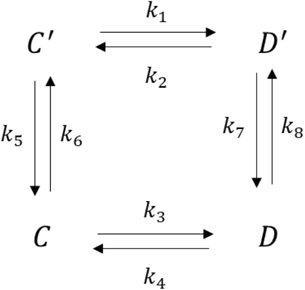

Here, C’, C, D’, and D denote the number of nucleosomes in these states, respectively. *K_1-8_* denote the transition rates between them, which will be defined below. Gene activation is defined to be the system state where all nucleosomes exist in a decompacted state. The two mechanisms are intertwined so that methylation states affect compaction rates and vice versa. Detailed descriptions of the rates are given below:

#### Methylation

In this model, un-methylated nucleosomes convert into a methylated state with a first-order rate constant β. We assume this rate constant is the same regardless of whether nucleosomes are inside or outside the compacted assembly. Methylated nucleosomes convert into a de-methylated state with a rate constant of *α* if the nucleosome is outside the assembly (D’), or a lower rate constant of *fα*, (*f* < 1) if the nucleosome is inside the assembly. This lower rate constant assumes that the demethylation reaction is less efficient on compacted nucleosomes, possibly due to competition for demethylase binding by compaction proteins, or due to the exclusion of demethylases through steric occlusion or phase separation. The rates describing these reactions on the four nucleosomal species are given by:

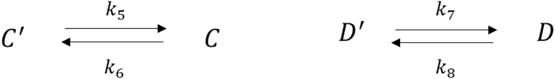

Where:

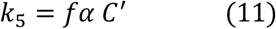

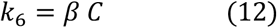

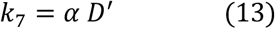

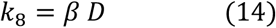

H3K27 methylation and demethylation rates are based on the catalytic activity of the EZH2 subunit the PRC2 complex and Kdm6a/b proteins, respectively. Specifically, these rate constants were chosen to represent the conversion between H3K27me2 and H3K27me3. For simplicity, we do not model the H3K27me-binding dependent H3K27 methylation activity previously described (Margueron et al., 2009), though we show below that explicit modeling of this cooperative effect would not significantly alter the conclusion of the model. JMJD3 and UTX demethylate H3K27me3 (Agger et al., 2007), and to our knowledge no cooperative activity of these complexes have not been reported.

#### Compaction

We adopt a mean-field description of the compacted nucleosomal assembly, following kinetic models of multi-stranded cytoskeletal polymer assembly(Howard, 2001). This description assumes that the nucleosome assembly is a roughly spherical structure held together by weak, multivalent interactions between individual nucleosomes, and can add or lose individual nucleosomes at its surface. Both methylated and demethylated nucleosomes can incorporate into the assembly; thus the assembly has a total size of:

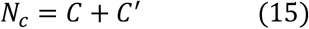

where *C* and *C*′ represent the number of methylated and demethylated nucleosomes in the assembly, respectively. Unlike other polymer models (MacPherson et al., 2018; Nuebler et al., 2018), we do not explicitly model physical connections between nucleosomes due to DNA; such connections would be expected to result in a spatial dependence of reaction rates within this chromatin domain; however, as the entire domain (100 nucleosomes) has a length scale greater than the persistence length of chromatin (∼15-20 nucleosomes, from (Arbona et al., 2017), and would thus enable free interactions between non-neighboring nucleosomes, we would expect the essential properties of our minimal model would also hold in a more realistic physical model that incorporates nucleosome connectedness.

The addition and removal of methylated and demethylated nucleosomes from the assembly is described by the following rate equations:

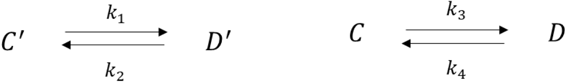

Where:

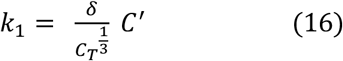

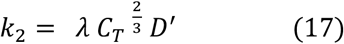

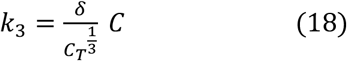

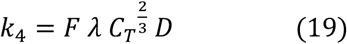

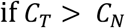

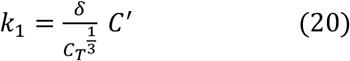

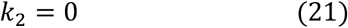

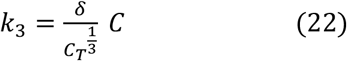

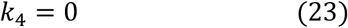

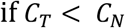

Here, methylated nucleosomes incorporate into the compacted assembly with a rate constant v; however, importantly, demethylated nucleosomes can also incorporate into the assembly with a reduced rate constant *Fλ* (where *F* < 1). The effect of methylation state on compaction rate is experimentally observed in instances such as recruitment of PRC1 complex by H3K27me3 marks (Kahn et al., 2016). The complex’s subunits such as Ring1B and Phc-1 have been shown to be important in chromatin compaction and gene silencing (Eskeland et al., 2010; Francis et al., 2004; Isono et al., 2013). However, as PRC1 recruitment is not the only compaction mechanism *in vivo*, and because PRC1 can bind to nucleosomes independently of H3K27me3 (Francis et al., 2004), this model treats methylation as only a part, but not solely responsible for chromatin condensation. In choosing rate constants; we assume that compaction and decompaction is faster than histone methylation and demethylation rates, though timescales for both processes are assumed to be much faster than that for cell division. Fast compaction kinetics relative of modification is supported by *in vitro* studies of H3K27me3 methylation and demethylation kinetics, as well as *in vitro* DNA compaction by HP1_α_ and chromatin condensation experiments (Kristensen et al., 2011; Ladoux et al., 2000; Larson et al., 2017; Sneeringer et al., 2010).

The reaction rates for nucleosome incorporation (loss) scales with assembly size as 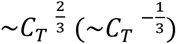, as these reactions only take place on the surface of the assembly. Assuming a compacted nucleosome complex is spherical, the compaction rate would thus be proportional to the surface area. Likewise, the decompaction rate is also proportional to the surface area but reversely proportional to the total number compacted nucleosome in the complex.

In this description, there is a critical threshold number of compacted nucleosomes, *C_N_* below which the complex is thermodynamically unstable. The existence of a minimal nucleus size is a fundamental property of multivalent polymers, whereby addition of a new subunit to an already formed complex is thermodynamically more favorable than formation of the initial nucleus itself (Erickson and Pantaloni, 1981; Jackson and Berkowitz, 1980). Below this critical threshold *C_N_*, the compacted assembly disintegrates, and gene turns on.

#### Cell division

Heritability of histone marks and chromatin states are crucial in maintaining gene expression states across cellular generations. As with the methylation only model above, we assume that methylated nucleosomes partition randomly between two daughter strands upon replication; the total number of nucleated nucleosomes is then obtained by sampling from binomial distribution with *p = 0.5* and *N* equal to the total number of nucleosomes at the point of DNA replication. Furthermore, we assume that compacted nucleosomes persist within a compacted assembly reside upon passage of DNA polymerase. This model feature assumes that new nucleosomes rapidly incorporate into a compacted assembly after passage of DNA polymerase; however, in the subsequent version of this model below, we will relax this assumption to allow for disruption of compaction state by DNA polymerase passage (see below).

From Monte-Carlo simulations, we found that this methylation compaction model can recapitulate all the essential emergent properties of the *Bcl11b* activation switch. Specifically, this model shows the following dynamic properties:

1. *Irreversible all-or-none switching to an H3K27me3-low, de-compacted state.* From simulations, we found that the system adopts a stable compacted assembly of nucleosomes with higher H3K27me3 marking density, but switches abruptly to a de-compacted state with lower H3K72me3 levels. As there is no re-nucleation of the compacted assembly after its elimination, this de-compacted state represents an absorbing, permanently active expressing state. The abrupt decrease in the H3K27me3 levels arises because compacted nucleosomes demethylate at a lower rate; thus, upon total decompaction, the percent of methylated nucleosomes lowers to a new steady state level.
2. *Noise induced gene activation.* Transition to the completely decompacted state, or gene activated state, occurs via stochastic deviation of the system from its compaction meta-stable state. Activation is triggered when the system reaches below the threshold number of compacted nucleosomes.
3. *Tunable activation rates*. The model is able to generate a gene switch with slow, tunable activation rate. Activation delays can span many days, and can be finely adjusted by modifying methylation and demethylation rates, and/or changing H3K27me3 levels at the gene locus (Fig. 4C,G,H), as experimentally observed (Fig. 3). This ability to tune activation rates by changing H3K27me3 densities uniquely distinguishes the methylation compaction model from the methylation only model above, and thus represents a more plausible model for describing the activation mechanism of this switch. Why is this model uniquely tunable? In this model, locus de-compaction and gene activation are determined by a dynamic balance between rates of nucleosome entry or exit from a compacted assembly. The system still be sensitive to changes in these rates; however, as demethylated nucleosomes can still enter and exit a compacted assembly at a reduced rate, changes in the fraction of demethylated nucleosomes would cause a fine change in these entry or exit rates, and thus represent a plausible tuning parameter for controlling activation timing.
4. *Division-independent timing control*. When the cell cycle length is changed in this model, activation kinetics remain largely unaffected, implying that the methylation-compaction mechanism functions as a cell division-independent delay timer. These conclusions hold, as long as the dynamic methylation and compaction mechanisms operate on timescales much faster than the cell cycle length.

Why is the methylation compaction tunable? To answer this question, we adopt an approach, where we reduce this problem using the Fokker-Planck approximation, as utilized to analyze the Methylation only mechanism (Fig. S10A). The full system with both methylation and compaction reactions would correspond to diffusive motion of a particle in a three-dimensional state space describing both chemical and physical states of nucleosomes. However, to simplify this problem to gain intuition, we will first take the methylation and demethylation reactions to be fast compared to the compaction and de-compaction reactions, such that the system can be described a single parameter N_C,_ corresponding to the total number of compacted nucleosomes. At any given time, the number of methylated and demethylated nucleosomes in the compacted state is at quasi-steady state, with values:

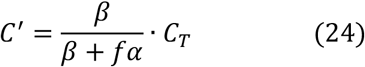

and

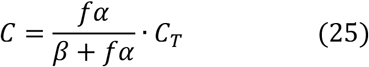

Similarly, assuming quasi steady-state, the number of methylated and demethylated nucleosomes in the uncompacted state is given by:

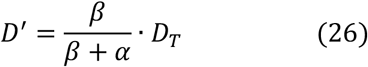

and

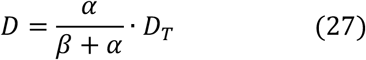

Let *N_T_*= *C_T_* + *D_T._* With this approximation, the averaged rate of adding or removing a nucleosome from the compacted assembly is then given by:

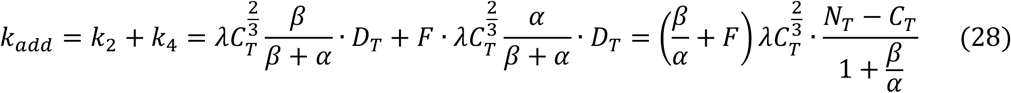

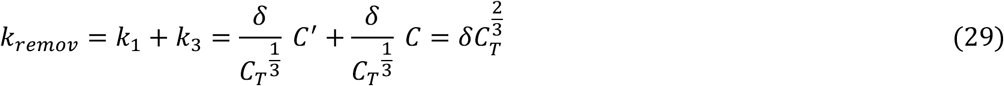

Let the total number of compacted nucleosomes *C_T_* be *x*. By writing down the master equation for this system, and by further applying the Fokker-Planck approximation, as performed in (5) and (6) we then have:

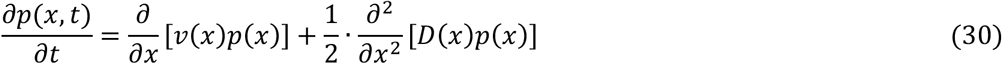

where:

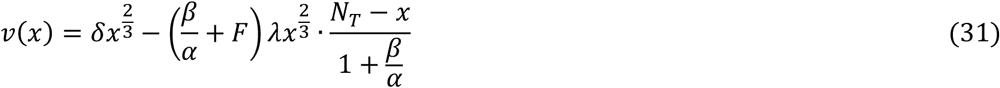

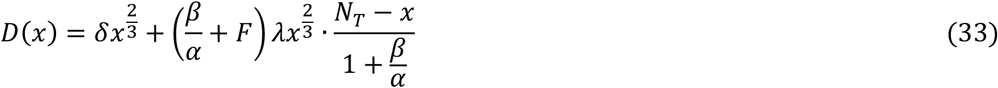

As before, we define a potential energy for this system:

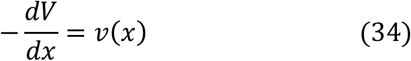

The analytical solution for the potential energy *V*(*x*) for the methylation compaction model is:

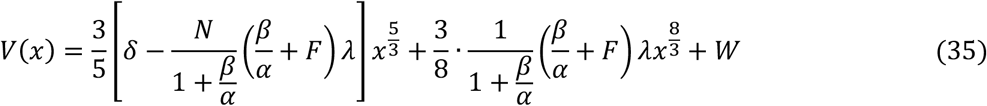

A plot of *V*(*x*) is shown in Fig. S10B-C. We found that increasing methylation rate results in a much more attenuated increase in activation energy *E_a_* with the methylation compaction model. This confirms that the improved switching rate tunability in the MC model stems from the decreased sensitivity to changes in activation barrier height by methylation rate. This result explains why this system shows significantly more graded changes in switching times when methylation rates are changed.

This tunability of switching times with respect to histone methylation depends on the relative association strengths of demethylated and methylated nucleosomes for each other in forming a compacted assembly. In our initial simulations, demethylated nucleosomes show only a moderate decrease in affinity for other compacted nucleosomes relative to methylated nucleosome (*F = 0.85*). However, when the binding strength of a demethylated nucleosome is much weaker than that of a methylated nucleosome (*F = 0.2*), we find changes in potential well heights become more significant, indicating that the system loses its tunability with respect to methylation changes (see Fig. S10B-C). This prediction, that demethylated nucleosomes maintain comparable strengths of self-association for compacted assembly formation, agrees well with evidence that unmethylated nucleosomes aggregate through a variety of H3K27me-independent mechanisms (Larson et al., 2017; Strom et al., 2017).

#### Model II.1: The Methylation Compaction Mechanism, with Compaction Disrupted by Division (**Fig. S2**)

This version of the model includes modified cellular division process in which upon replication, 50% of methylated nucleosomes become demethylated and 10% of compacted nucleosomes become uncompacted. This exit of nucleosomes from a compacted assembly due to DNA replication reflects the possibility that as the DNA replication machinery enters the compacted nucleosomal structure, it creates decompaction ‘defects’ in the condensed locus because nucleosomes near the replication forks are replaced. However, we reason that such defect would have a small effect to the overall stability of the structure because, at any given time, the site of replication would only take up a small region of the entire compacted domain.

In order to simulate both changes in methylation and compaction at the point of DNA replication, we must describe probabilistically how each of the four nucleosomal species are affected: 1) The Compacted-Methylated species (C’); 2) the Compacted-Demethylated species (C); 3) the Decompacted-Methylated species (D’); and 4) the Compacted-Demethylated species (D). Since methylation state is reduced by 50%, approximately half of Decompacted-Methylated species is transferred to Decompacted-Demethylated pool. Similarly, on average, 10% of the Compacted-Demethylated species are transferred to Decompacted-Demethylated pool due to DNA replication. Compacted-Methylated species have 50% chance to demethylate and 10% chance to decompact. Assuming these are two independent processes, this species has 5% chance to convert into Decompacted-Demethylated or Decompacted-Methylated and 45% chance to become Compacted-Demethylated. These observations are implemented as follows:

Let vector ***S*** = [*S*_1_, *S*_2_, *S*_3_, …, *S_n_*] be the result from sampling a multinomial distribution with probabilities *π*_1_, *π*_2_, *π*_3_, … , *π_n_*, where *π*_1_ + *π*_2_ + *π*_3_ + ⋯ + *π_n_* = 1. Let *S_i_*(N|*π*_1_, *π*_2_, *π*_3_, … , *π_n_*) be the *i*^th^ element of ***S*** and N be the sample size. Let *c’*, *c’*, *d’*, b be the number of compacted-methylated, compacted-unmethylated, decompacted-methylated, and decompacted-unmethylated nucleosomes, respectively immediately preceding the cellular division event. Partitioning of each species occurs as follows:

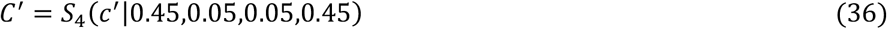

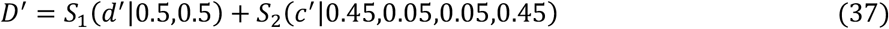

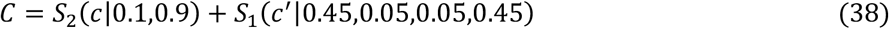

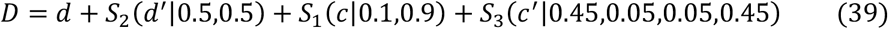

From stochastic simulations (Fig. S2), we find that this modified methylation compaction model shows similar dynamic characteristics compared to the original methylation compaction model (Model II): it shows stochastic, all-or-none switching between inactive and active states; has an activation delay that can be tuned by changing H3K27me levels and enzyme activity; and shows division-independence in its activation time delay. Thus, we conclude that the essential features of this model hold, even upon mild disruption of the inactive, compacted assembly by passage of DNA polymerase.

#### Model II.2: The Methylation Compaction Mechanism with Cooperative Methylation (**Fig. S3**)

PRC2 is known to be allosterically activated by H3K27me3 binding via its EED subunit (Margueron et al., 2009). Here, we consider this cooperative property of PRC2 by specifying that methylation rate increases with the total number of methylated nucleosomes in the model system. This assumption is likely valid when the number of nucleosomes in the condensed structure is small, and all the nucleosomes are more or less in close proximity with each other. To simulate this, we modified the methylation rates *K*_6_ and *K*_8_ so that their magnitude has a spontaneous term *μ* and the cooperative term *β* that is proportional to the total number of methylated species in the simulation:

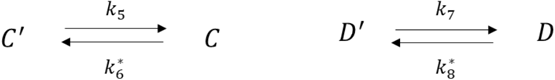

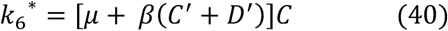

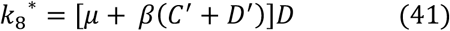

From stochastic simulations (Fig. S3), we find that this system is also capable of generating long, stochastic delays in all-or-none switching in locus compaction state, and that switching times can be finely tuned by changing H3K27me3 levels, as with our simpler methylation compaction model (Model II). We conclude that incorporation of a cooperative H3K27me3 methylation rate in our methylation compaction model does not alter its main conclusions.

#### Model II.3: The Methylation Compaction Mechanism with Permanently Demethylated Nucleosomes (**Fig. S4**)

To simulate the potential effect of transcription factors on the activation timing in the methylation compaction model. We first consider a scenario where the transcription factors disable a small fraction of nucleosomes in the assembly from compacting or methylating, effectively removing them from the system. Our simulations (Fig. S4A,B) suggests that activation timing is faster with increasing number of removed nucleosome (*N_r_*). Therefore, timing modulation can be achieved this way although activation time is mildly sensitive to changes in *N_r_*.

We then consider an alternative mechanism where the transcription factors act on the system by binding to a small number of nucleosomes and preventing it from being methylated but still allowing compaction. To formulate this effect, we define a sub population of nucleosomes in the system that can undergo compaction *C*_o_ and decompaction *D_O_* but cannot be methylated:

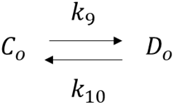

Where:

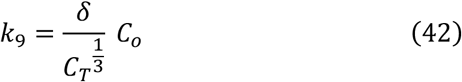

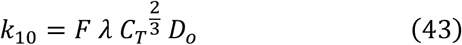

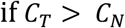

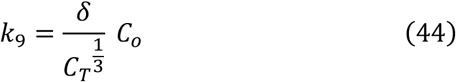

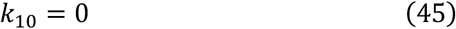

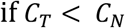

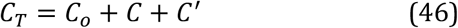

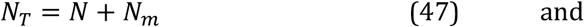

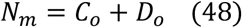

Simulations of this system (Fig. S4C) reveals that activation timing is faster with increasing number of permanently demethylated nucleosomes. Additionally, the changes in timing is not as drastic as completely removing nucleosome from the system altogether. These results present potential mechanisms through which transcription factors could be employed to manipulate activation timing in polycomb regulated gene loci.

### Part II: Modeling a regulatory gene circuit composed of epigenetic timers

#### Introduction

During development, progenitors maintain autonomous temporal schedules for differentiation to ensure proper cell expansion. These temporal schedules are precisely specified within an organism but can be scaled across species to vary organism sizes while maintaining basic tissue proportions. Here, we ask whether regulatory gene circuits built from individual timed epigenetic switches can implement temporal schedules for differentiation that are robustly defined, yet flexibly scalable, such the expression timing between each node can be modulated over a large dynamic range. We consider a simple regulatory circuit consisting of a series of genes activating in sequence, where each gene is turned on by an epigenetic timer, and in turn activates an epigenetic timer of a downstream gene. As a performance comparison, we also consider a second gene network, where genes also activate each other in sequence, but do so through classical gene regulation functions, where transcription rates are functions of the levels of upstream regulator (Estrada et al., 2016; Phillips, 2015). We model these networks at the single-cell level, performing Monte-Carlo simulations of network dynamics in single cells, were we explicitly model cell division. To analyze variability in the dynamics of these networks, both in single cells and in cohorts of developing cells, we perform this analysis over multiple cohorts of developing cell populations.

#### Model I: Regulatory network of epigenetic timer

In our model, each cell contains a simple sequential gene network, where an initiating factor *X*_0_ initiates the sequential activation of a series of genes *X*_1_, *X*_1_, *X*_3_, and *X*_4_ In this cascade, an upstream transcription factor *X*_*i*-1_ activates a epigenetic timer for a single promoter *p_i_* (*i* = 1, …, 4), which sets the activation time for synthesis of its product *X_i_*. . The control of activation follows this scheme:

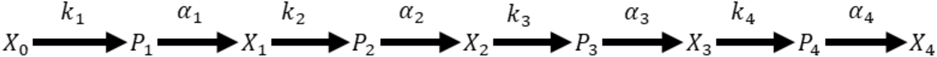

For simplicity, we assume that each cell has one copy of each gene, such that *p_i_* is a random variable with two possible states (0 and 1). However, we expect our main conclusions to hold when multiple gene copies are present. *X_i_* is then a random variable that represents the copy number of the product of gene *i* in the cell.

Take *K_i_* to be the rate constant for activation of promoter *i*. We take *K_i_* to be hyperbolic function of the concentration of upstream regulatory factor, such that:

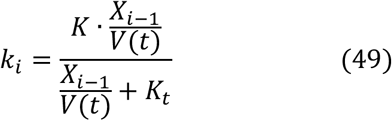

where *K* denotes the maximal rate of promoter activation, *K_t_* is the concentration of transcription factor required for half-maximal promoter activation rate, and *V*(*t*) is the volume of the cell, which varies as a function of the cell cycle. Biologically*, X_i_* could promote activation through a variety of possible mechanisms, including recruitment of H3K27me3 demethylases, inhibition of PRC2 methyltransferase activity, or any other mechanisms that would lead to a local disruption of H3K72me3 distributions in the vicinity of transcription factor binding. In accordance with the observed properties of the *Bcl11b* epigenetic timer, we take the maximal activation rate to be less than that of cell division, and also take this rate to be independent of the rate of cell division itself, which we independently modulate below.

In addition to being modulated by specific transcription factors, H3K27me3 levels at gene loci – and, by extension, gene activation rates – can also be tuned by global activities of H3K27me3-modifying enzymes. Specifically, cells with lower H3K27me3 methylation activity will have gene loci with a higher maximal activation rate, due to low basal repressive mark level. We do not explicitly model levels of H3K27 modifying enzyme in the cell, but model effects of their change through a coordinated change in the maximal rate of gene activation *K*. We take the value of *K* to be the same for all genes in the network, and assume that changes in H3K27 modifying enzyme activity will have a coordinated effect on all loci. We define a delay parameter, *d_E_*, as follows:

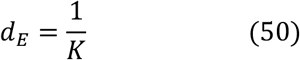

The synthesis and degradation of transcription factors å = 1 … 4 occur with first order kinetics, with reactions and rates given by:

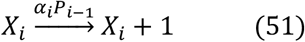

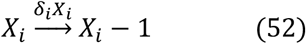

For the first equation, it can be seen that synthesis occurs only when the promoter is active. For simplicity, we set transcription rate α*_i_* and degradation rate *δ*_i_ to be the same for all species, and we assume these rates remain fixed throughout cell cycle.

During cell division,*X_i_* is binomially partitioned with a probability of 50% between two daughter cells upon cell division, and also inactive and active states of their promoters are stably inherited, consistent with heritable nature of the *Bcl11b* allelic expression states observed. We also assume that the level of the starting signaling molecule *X*_0_ remains constant, though we note that persistent signals are not required to maintain the states of the transcription factors after activation.

Finally, we adopt a discrete modeling approach to simulate each cell cycle separately. Taking n = 1…*N* to be the total number of cell cycles, *T* to be the total number of simulated timesteps within one cycle, and τ to be the total simulation time, we calculate *V*(τ) is as follows:

Simulation time τ = (*n* + *t*)Δ*t* (Δ*t* « 0)

For *n* = 1,2,3, …, *N*:

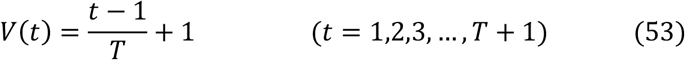

such that the cell cycle length is *T*ΔV. Note that *V* increases linearly between 1 and 2 at every cell cycle, and that the maximum and minimum cell volumes are not affected how the speed of cell division.

From stochastic simulations, we found that this network can give rise to a defined temporal schedule for differentiation within cohorts of developing cells, with a timescale exceeding that of cell division (every 20 hours). Furthermore, this schedule can be concurrently lengthened or shortened by changing the global delay parameter (Fig. 6B-C). Interestingly, the timing of gene activation, defined as the time it takes for concentration of *Xi* to reach a given threshold level, is independent of the rate of cell division (Fig. S9A-C). These results indicate that polycomb-mediated epigenetic timers can be utilized to generate scalable temporal delays in the differentiation schedules of progenitor cells.

#### Model II: Classical gene regulation network

The timing of gene expression can also be set in gene networks through use of classical gene regulation functions, where the rate of transcription is a function of the levels of upstream transcription factor. Here, we tested whether such classical regulatory gene networks are similarly capable of generating scalable differentiation schedules. We simulated a series of genes sequentially connected to each other, where each preceding gene acts as a transcriptional activator for the next gene:

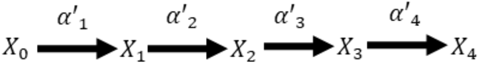

We reasoned that if interaction between transcription factor activator and the promoter is weak, it will take longer for the upstream species to accumulate to a sufficient threshold level to activate of the downstream species. The rate of transcription of gene X_i_ is given by:

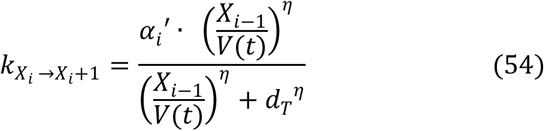

Here, *α_i_*’ is the maximum transcription rate achieved by the transcription activator, and *d_T_* is the dissociation constant of the activator to the promoter. We set *α_i_*’ to be the same for all species. We modeled the activation as a Hill Function with high cooperativity η to create a sharp threshold for switching. With this setup, gene expression of downstream species does not begin until the concentration of the activator reaches *d_T_*, such that that increasing this parameter would ultimately lead to lengthening of activation delay. Cell volume and species’ copy number dynamics as a function of cell cycle remain the same as in Model I. However, we assume that *X_i_* is a stable proteins that only becomes removed through cell-cycle mediated dilution. This stability is required for generating differentiation schedules with timescales that exceed that of the cell division time.

Stochastic simulation of this model revealed that multi-generational delay in gene expression can be achieved with this gene network architecture (Fig. 6F, G). However, this delay is non-scalable as increasing *d_T_* pass the steady state concentration of the upstream gene species will eventually result in a failure to activate downstream genes (Fig. 6G). As a result, activation delays set by this this network cannot be tuned beyond a limited range. Additionally, since protein degradation is tied to cell cycle division, both timing of and amplitude of gene expression are under influence of cycling time (Fig. S8D-F). Particularly, gene expression amplitude increases with lengthening cell cycle; this in turn speeds up activation of downstream species due increase in signaling molecules. In conclusion, networks built using classical gene regulation functions can set schedules over short timescales; it does not operate well over long timescales, and cannot be scaled in time through adjustment of a global transcriptional parameter.

In summary, we find that gene regulatory networks built from individual polycomb-mediated epigenetic timers can generate robust temporal schedules for differentiation that are tunable through the control of a single parameter, namely, the strength of epigenetic repression. This network uncouples timing from both cellular division and expression magnitude. In contrast, gene regulatory networks built using classical gene regulation functions cannot generate schedules over long times, and do not facilitate scaling of these schedules independently from cell division.

#### Model for progenitor differentiation dynamics

To understand whether the epigenetic timer cascade can give rise to proportional changes in the total number of output cells of different types, we integrated the gene regulatory network models above with a differentiation scheme, where progenitors generate different cell types in a progressive manner through asymmetric cell division. This scheme follows the general model described for temporal neuronal specification in flies and mammals (Rossi et al., 2017).

At every cell division, a single progenitor divides to give rise to a copy of itself and a terminally differentiated cell. The differentiated cell type is specified by the regulatory gene that has most recently activated and reached a threshold (Figure 6A). The timing to reach threshold for each gene species is extracted from gene network simulations described in the previous sections (Model I and Model II). Let *T_Xi_*be the time at which gene species *X_i_* reaches a threshold, *t_k_* be the time of *k*_th_ cell division event, and *C_i_* be the total number of differentiated cell type *i* The algorithm for such an asymmetrical differentiation scheme is as follows:

*For k = 1, 2, 3, …, n:*

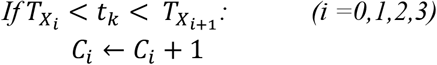

This process of asymmetric division continues until the progenitor turns on *X*_4_, which causes the progenitors to stop dividing and the simulation to terminate. From our simulations, we found that timing delays generated by networks of epigenetic timer give rise scalable population size of each differentiated cell type over a wide range of delay parameters (*d_E_*) (Fig. 6D). Furthermore, the coefficient of variation in the numbers of different cells produced per cohort decreased steadily with increasing cohort size, indicating that this system can precisely specify the proportions of output cells with sufficient initial progenitor numbers (Fig. 6F). On the other hand, the classical regulation network lead to severely disproportionate expansion of early cells in the cascade upon increase in the timing delay parameter (*d_T_*) (Fig. 6H). This selective expansion occurred as later regulatory genes failed to activate, causing the arrest of earlier progenitors in a proliferative state. In summary, regulatory networks built from epigenetic timers are uniquely suited for scaling of temporal schedules and population sizes through adjustment timing control parameters.

### Parameter List

#### a. Pure Methylation Model (Figure 4B-C)

| Pure Methylation Model |  |  |  |
| --- | --- | --- | --- |
| Parameters | Description | Value | Comments/references |
| $\beta_{ON}$ | cooperative methylation rate constant | 1 hrs<sup>-1</sup> | Sneeringer et al., 2010 |
| $\alpha_{OFF}$ | demethylation rate constant | 16.5 hrs<sup>-1</sup> | [16, 17, 17.5] for parameter scan<br>Kristensen et al., 2011 |
| L | reach of anchored methylating enzyme to neighboring nucleosome | 15 | Hass et al., 2015 |
| N | number of simulated nucleosomes | 100 |  |
| cell division length | cell division length | 20 hrs |  |

#### b. Compaction Methylation Model (Figure 4C,E)

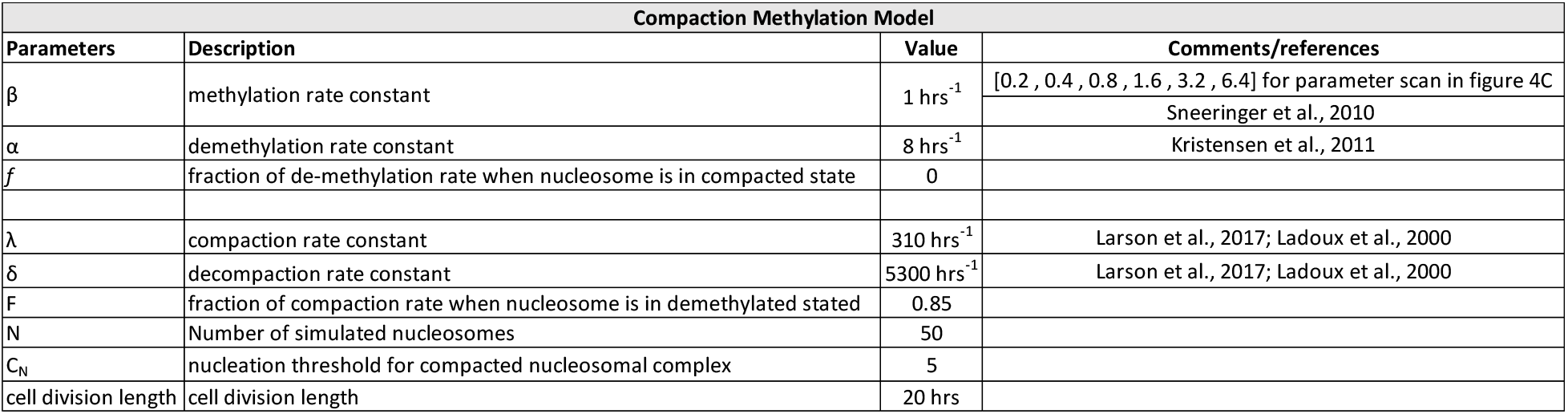

| Compaction Methylation Model |  |  |  |
| --- | --- | --- | --- |
| Parameters | Description | Value | Comments/references |
| $\beta$ | methylation rate constant | 1 hrs<sup>-1</sup> | [0.2, 0.4, 0.8, 1.6, 3.2, 6.4] for parameter scan in figure 4C<br>Sneeringer et al., 2010 |
| $\alpha$ | demethylation rate constant | 8 hrs<sup>-1</sup> | Kristensen et al., 2011 |
| f | fraction of de-methylation rate when nucleosome is in compacted state | 0 |  |
| $\lambda$ | compaction rate constant | 310 hrs<sup>-1</sup> | Larson et al., 2017; Ladoux et al., 2000 |
| $\delta$ | decompaction rate constant | 5300 hrs<sup>-1</sup> | Larson et al., 2017; Ladoux et al., 2000 |
| F | fraction of compaction rate when nucleosome is in demethylated stated | 0.85 |  |
| N | Number of simulated nucleosomes | 50 |  |
| C<sub>N</sub> | nucleation threshold for compacted nucleosomal complex | 5 |  |
| cell division length | cell division length | 20 hrs |  |

#### c. Pure Dilution and Noise vs induced Decompaction Models (Figure 5 A-B)

| Pure Dilution Model |  |  |  |
| --- | --- | --- | --- |
| Parameters | Description | Value | Comments/references |
| $\beta_{ON}$ | cooperative methylation rate constant | 0 | |
| $\alpha_{OFF}$ | demethylation rate constant | 0 | |
| L | reach of anchored methylating enzyme to neighboring nucleosome | 0 |  |
| N | number of simulated nucleosomes | 100 |  |
| cell division length | cell division length | 20 hrs | [10, 15, 20, 25, 30, 35] for cycle scan in Figure 5B |
| Compaction Methylation Model |  |  |  |
| Parameters | Description | Value | Comments/references |
| $\beta$ | methylation rate constant | 1 hrs<sup>-1</sup> | Sneeringer et al., 2010 |
| $\alpha$ | demethylation rate constant | 8 hrs<sup>-1</sup> | Kristensen et al., 2011 |
| f | fraction of de-methylation rate when nucleosome is in compacted state | 0 |  |
| $\lambda$ | compaction rate constant | 310 hrs<sup>-1</sup> | |
| $\delta$ | decompaction rate constant | 5300 hrs<sup>-1</sup> | |
| F | fraction of compaction rate when nucleosome is in demethylated stated | 0.85 |  |
| N | Number of simulated nucleosomes | 50 |  |
| C<sub>N</sub> | nucleation threshold for compacted nucleosomal complex | 5 |  |
| cell division length | cell division length | 20 hrs | [10, 15, 20, 25, 30, 35] for cycle scan in Figure 5B |

#### d. Polycomb-based timer model and classical regulation timer model (Figure 6)

| Polycomb-based timer model |  |  |  |
| --- | --- | --- | --- |
| Parameters | Description | Value | Comments/references |
| d<sub>E</sub> | global epigenetic switching rate delay | 25 and 66.66 hrs | [100, 66.67, 50, 40, 33.33, 28.57, 25, 22.22, 20] for parameter scan |
| kt | dissociation constant of polycomb transcription factor | 400 copies / cell volume |  |
| $\alpha$ | transcriptional rate | 1000 copies hrs<sup>-1</sup> | Schwanhäusser, B. et al., 2011 |
| $\delta$ | degradation rate | 1 copies hrs<sup>-1</sup> | Schwanhäusser, B. et al., 2011 |
| cell division length | cell division length | 20 hrs |  |
| Classical regulation timer model |  |  |  |
| Parameters | Description | Value | Comments/references |
| d<sub>T</sub> | dissociation constant of transcription factor | 150 and 200 copies / cell volume | [20, 40, 60, 80, 100, 120, 140, 160, 175] for parameter scan |
| $\alpha'$ | transcription rate constant | 10 copies hrs<sup>-1</sup> | Schwanhäusser, B. et al., 2011 |
| $\eta$ | Hill coefficient | 50 | |
| cell division length | cell division length | 20 hrs |  |

#### e. Compaction Methylation Model with Compaction State Affected by Cell Division(Supplementary Figure 2)

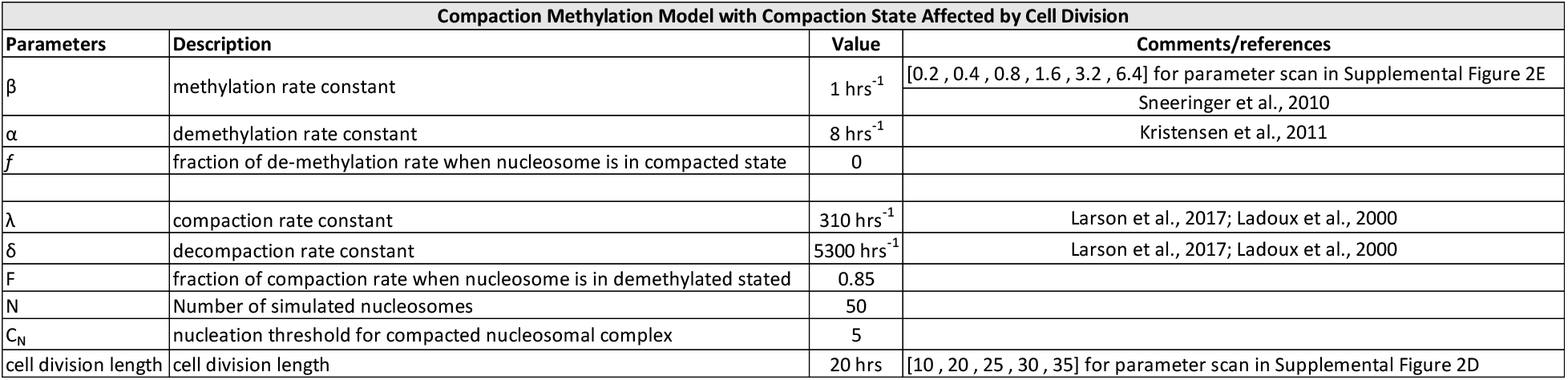

#### f. Compaction with Cooperative Methylation Model (Supplementary Figure 3)

| Compaction Cooperative Methylation Model |  |  |  |
| --- | --- | --- | --- |
| Parameters | Description | Value | Comments/references |
| $\beta$ | cooperative methylation rate constant | 0.02 hrs<sup>-1</sup> | |
| $\mu$ | spontaneous rate constant | [0.05, 0.1, 0.2, 0.4] hrs<sup>-1</sup> | |
| $\alpha$ | demethylation rate constant | 8 hrs<sup>-1</sup> | Kristensen et al., 2011 |
| $f$ | fraction of de-methylation rate when nucleosome is in compacted state | 0 | |
| $\lambda$ | compaction rate constant | 310 hrs<sup>-1</sup> | Larson et al., 2017; Ladoux et al., 2000 |
| $\delta$ | decompaction rate constant | 5300 hrs<sup>-1</sup> | Larson et al., 2017; Ladoux et al., 2000 |
| $F$ | fraction of compaction rate when nucleosome is in demethylated stated | 0.85 | |
| $N$ | number of simulated nucleosomes | 50 | |
| $C_N$ | nucleation threshold for compacted nucleosomal complex | 5 | |
| cell division length | cell division length | 20 hrs |  |
| Pure Methylation Model |  |  |  |
| Parameters | Description | Value | Comments |
| $\beta_{ON}$ | cooperative methylation rate constant | 1 hrs<sup>-1</sup> | Sneeringer et al., 2010 |
| $\alpha_{OFF}$ | demethylation rate constant | [16, 17, 17.5] hrs<sup>-1</sup> | for parameter scan, Kristensen et al., 2011 |
| $L$ | reach of anchored methylating enzyme to neighboring nucleosome | 15 | Hass et al., 2015 |
| $N$ | number of simulated nucleosomes | 100 | |
| cell division length | cell division length | 20 hrs |  |

#### g. Effects of Transcription Factors on Methylation Compaction Model’s Activation Timing (Supplemental Figure 4)

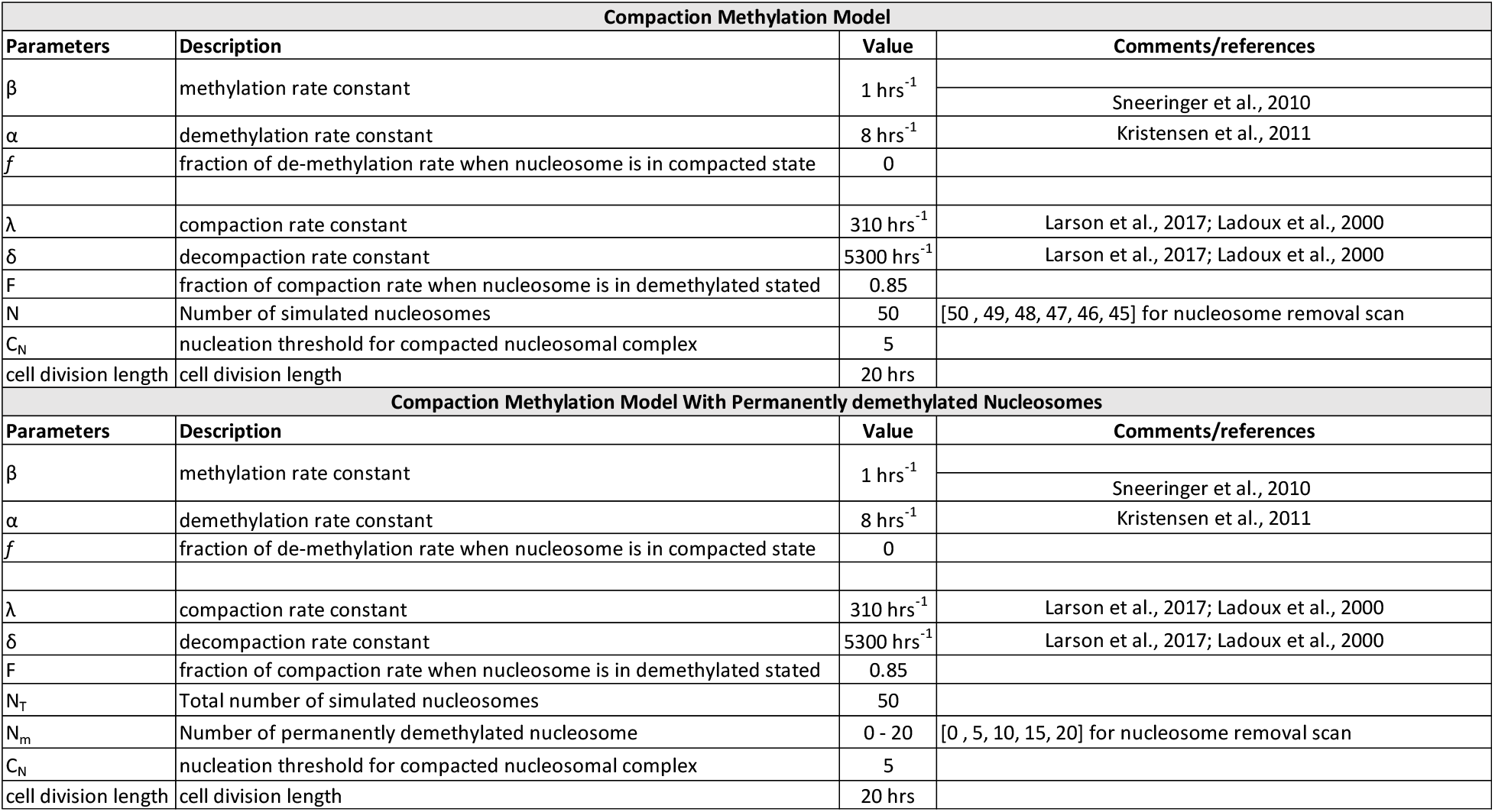

#### h. Pure Dilution Model with Minimal Methylation and Demethylation Rates (Supplementary Figure 5)

| Dilution Model With Minimal Enzymatic Activities |  |  |  |
| --- | --- | --- | --- |
| Parameters | Description | Value | Comments/references |
| $\beta_{\text{ON}}$ | cooperative methylation rate constant | 0.001 hrs<sup>-1</sup> | |
| $\alpha_{\text{OFF}}$ | demethylation rate constant | 0.001 hrs<sup>-1</sup> | |
| L | reach of anchored methylating enzyme to neighboring nucleosome | 15 | Hass et al., 2015 |
| N | number of simulated nucleosomes | 100 |  |
| cell division length | cell division length | 10 and 20 hrs |  |

#### i. Cell cycle dependency of Polycomb-based timer model and Classical regulation timer model (Supplementary Figure 9)

| Polycomb-based timer model |  |  |  |
| --- | --- | --- | --- |
| Parameters | Description | Value | Comments/references |
| $d_E$ | global epigenetic switching rate delay | 50 hrs | |
| kt | dissociation constant of polycomb transcription factor | 400 copies / cell volume |  |
| $\alpha$ | transcriptional rate | 1000 copies hrs<sup>-1</sup> | Schwanhäusser, B. et al., 2011 |
| $\delta$ | degradation rate | 1 copies hrs<sup>-1</sup> | Schwanhäusser, B. et al., 2011 |
| cell division length | cell division length | 20 and 40 hrs | [10, 15, 20, 25, 30, 35, 40, 45, 50, 55] for cycle scan |
| classical regulation timer model |  |  |  |
| Parameters | Description | Value | Comments/references |
| $k'$ | dissociation constant of transcription factor | 150 copies / cell volume | |
| $\alpha'$ | transcription rate constant | 10 copies hrs<sup>-1</sup> | Schwanhäusser, B. et al., 2011 |
| $\eta$ | Hill coefficient | 50 | |
| cell division length | cell division length | 20 and 40 hrs | [20, 25, 30, 35, 40, 45, 50, 55] for cycle scan |

#### j. Potential energy landscapes analysis for pure methylation model and methylation compaction model (Supplementary Figure 10)

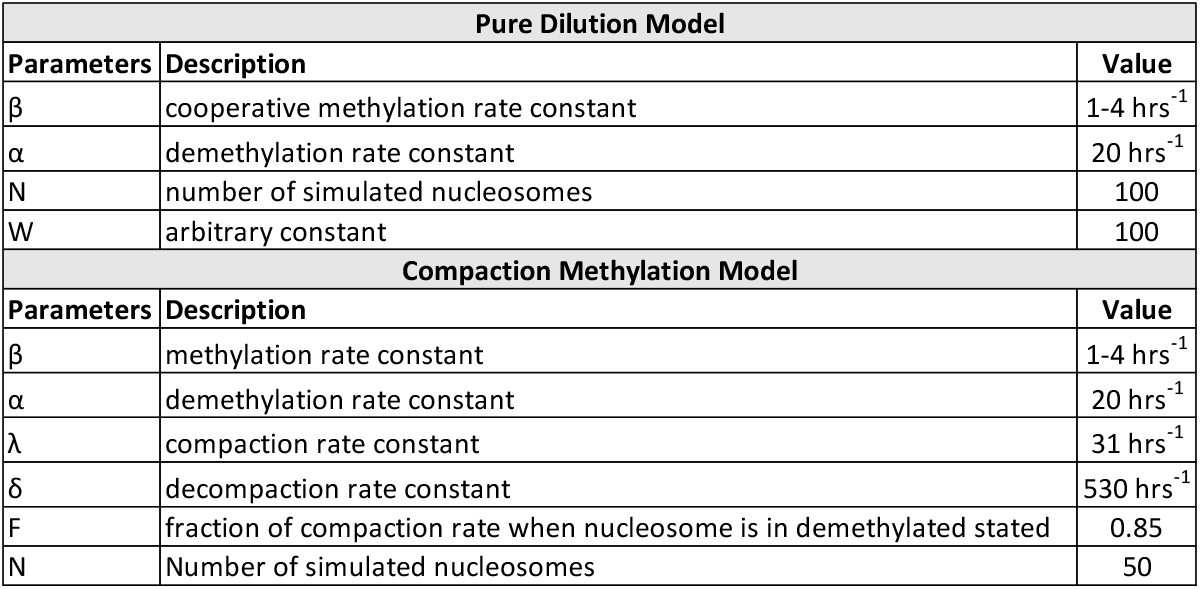

| Pure Dilution Model |  |  |
| --- | --- | --- |
| Parameters | Description | Value |
| $\beta$ | cooperative methylation rate constant | 1-4 hrs<sup>-1</sup> |
| $\alpha$ | demethylation rate constant | 20 hrs<sup>-1</sup> |
| N | number of simulated nucleosomes | 100 |
| W | arbitrary constant | 100 |
| Compaction Methylation Model |  |  |
| Parameters | Description | Value |
| $\beta$ | methylation rate constant | 1-4 hrs<sup>-1</sup> |
| $\alpha$ | demethylation rate constant | 20 hrs<sup>-1</sup> |
| $\lambda$ | compaction rate constant | 31 hrs<sup>-1</sup> |
| $\delta$ | decompaction rate constant | 530 hrs<sup>-1</sup> |
| F | fraction of compaction rate when nucleosome is in demethylated stated | 0.85 |
| N | Number of simulated nucleosomes | 50 |

